# Cerebral Cortical-Like Organoid Model of PPP2R5D Induced Genetic Intellectual Disability Displays Variant-Specific Disease Severity Phenotype

**DOI:** 10.64898/2026.05.26.728012

**Authors:** YuXin Du, Mandeep Singh, Mandar Patil, Isaac Villegas, Adriana Portillo, Kyle Shang, Roy Ben-Shalom, Julian Halmai, Kyle Fink

## Abstract

Jordan’s Syndrome (JS) is a rare, neurodevelopmental disorder caused by de novo missense mutations in protein phosphatase 2 regulatory subunit B’delta (*PPP2R5D*). JS is characterized by severe neurological impairments starting in early life. *PPP2R5D* encodes for B56δ, one of the regulatory subunits of protein phosphatase 2A (PP2A). PP2A is a heterotrimeric protein serine/threonine phosphatase that is highly expressed in the brain and the liver. Past studies have focused on PP2A’s role in liver and little is known about the holoenzyme’s behavior in neuronal cells. Although B56δ is known to play an important role in the substrate specificity of PP2A, the identification of validated downstream substrates in JS remains unclear. To better understand how the mutations affect neuronal cells, we developed cerebral cortical-like organoids from an engineered allele series of the most common JS mutations to characterize the physiological changes throughout different stages of neurodevelopment. Organoids were assessed for transcriptomic, protein, and electrophysiological changes utilizing bulk RNA sequencing, immunocytochemistry, Western Blot, and high-density MicroElectrode Array. The results identify differentially expressed genes and translated proteins, potential neuronal substrates, and significant electrophysiological signatures that suggest mutations in B56δ lead to variant-specific dysfunction of PP2A. Overexpression of *PPP2R5D* through AAV transduction of organoids rescued several phenotypes in the variants, suggesting different pathogenetic etiology underneath. Our findings successfully characterized cerebral cortical-like organoids in JS cell lines and demonstrated its potential as a model for studying neurodevelopmental disorder and for screening therapeutic approaches.

## Introduction

Houge-Janssens Syndrome 1^39^ (OMIM 616355), also known as Jordan’s Syndrome (JS), is a rare, neurodevelopmental disorder with early infant onset^1^, characterized by severe neurological impairments such as macrocephaly, frontal bossing, epilepsy, hypotonia, speech delay, gross motor delay, and behavioral challenges^1–3^. The disorder was first described in 2015^9^ and as of February 2026, there were more than 485 diagnosed cases in more than 50 countries. It is estimated that there are approximately 250,000 undiagnosed cases worldwide^8^. The genetic etiology of JS lies within the *de novo* heterozygous missense mutations in protein phosphatase 2 regulatory subunit B’delta (*PPP2R5D*)^1–6^ (OMIM 601646). *PPP2R5D* encodes for B56δ, one of the regulatory subunits of protein phosphatase 2A (PP2A)^12^. PP2A is a heterotrimeric protein serine/threonine phosphatase highly expressed in the brain and in the liver, primarily in neurons^12^, and is involved in many cellular processes^3^. Mutations in *PPP2R5D* are known to play an important role in the substrate specificity of the holoenzyme^3, 9–11^. Previous studies have shown PP2A-B56δ’s role as an inhibitor in the PI3K/AKT-mTOR signaling pathway^3, 9, 13^, demonstrating its importance in central nervous system development. Although JS is caused by a broad mutational spectrum, the variants E198K and E200K are the most common and E198K mutation alone makes up approximately 40% of the reported cases^31^. Generally, E198K patients displayed the most severe symptoms with more cognitive impairment^5^, less expressive language skills^5^, and more seizures (60.6% of patient population) ^5, 34^, while E200K patients display a milder phenotype^5, 27^. Additionally, it has been reported that E198K and E200K patients have been diagnosed with early-onset Parkinsonism^4^. The wide range of phenotypic behaviors of the dysfunctional protein indicate the disease as a spectrum disorder.

Past studies used non-human neuronal models (e.g., mouse models, HEK cells) and have been focused on the holoenzyme’s role in the liver, and little is known about PP2A-B56δ’s role in neuronal cells. In addition, although B56δ is known to play an important role in the substrate specificity of PP2A^3, 9–11^, the identification of validated downstream substrates in JS remains unclear. To investigate *PPP2R5D*’s role in human neuronal context, Carter et al^4^ and Young et al^6^ developed 2D neuronal culture as a model and identified genes and pathways associated with upregulation of excitatory neurons in the mutants. However, these models do not provide information regarding the changes in the trajectory of development as well as functional readouts of the neurons. To better understand how the mutations affect neuronal cell physiology across developmental stages and in a complex 3D environment, we developed cerebral cortical-like organoids from an engineered allele series of JS mutations . The organoids were assessed for transcriptomic, functional protein substrates, and electrophysiological changes utilizing RNA sequencing, immunocytochemistry (ICC), Western b lot, protein phosphorylation array, and high-density MicroElectrode Array (HD-MEA). The results identify differentially expressed genes and impacted molecular and cellular pathways, potential neuronal substrates, and electrophysiological phenotypes related to genetic variation and disease severity, providing characterization on the global phenotypes in the cerebral cortical context. Furthermore, due to previous reports on PP2A as a counter regulator on S6 kinase activities in Drosophila^15, 16^ and in HEK cells^40^, we tested whether Rapamycin could restore molecular and cellular phenotypes of JS mutations. Dosing with Rapamycin resulted in rescue of dysregulated neuronal firing, suggesting the drug’s therapeutic potential with further pre-clinical investigation. Finally, overexpression of healthy *PPP2R5D* showed rescued electrophysiological parameters in E198 and E200K, suggesting different pathogenetic etiology between the different variants. Our findings successfully characterized cerebral cortical-like organoids in JS cell lines and demonstrated its strong potential as a model for studying neurodevelopmental disorder and for screening therapeutic approaches.

## Results

### Generation of cerebral cortical-like organoids from iPSCs of PGP-1 allelic series

PGP-1 induced pluripotent stem cells (iPSCs) were engineered using CRISPR-HDR to generate an allelic series of mutations in *PPP2R5D*. PGP-1 Healthy, E198K, and E200K iPSCs were used in this study as they represent two common mutations with different clinical presentations. Sanger sequencing confirmed the heterozygous missense mutations at E198K and E200K sites in *PPP2R5D* (**Figure 1A**). Pluripotency was confirmed in the iPSCs using PODXL5 (**Figure 1C**). iPSCs were differentiated into neural stem cells (NSCs) with dual-SMAD inhibition. NSCs were positive for Nestin (**Figure 1D**) and were aggregated into cerebral cortical-like organoids (COs) using a three-phase process^32^ (**Figure 1B**). The development of COs was monitored through a panel of neuronal markers on the transcriptomic and histologic level (**Supplement Figure 1**). Quantitative real-time quantitative PCR (RT-qPCR) panel shows appearances of *MAP2* and *NeuN* on day 18 (d18) and increases over time on d46 and d90. Complemented with ICC results, these results demonstrate the cerebral cortical-like nature of the organoids. D123 and d184 ICC (**Figure 1E**) results show positivity in layer markers (SATB2, CTIP2), inhibitory marker (GAD67), excitatory marker (vGLUT1), and glial markers (S100B, GFAP). Throughout the differentiation process, we observed increased growth rate of E198K and E200K compared to Healthy, leading to the distinct size differences between the COs where E198K appeared to be the largest and Healthy were the smallest among the group (**Figure 1F**). Quantification of CO sizes shows significant differences between Healthy, E198K, and E200K on d18 (*p* < 0.001 for E198K, *p* = 0.0046 for E200K), d46 (*p* < 0.0001), and d90 (*p* < 0.0001). The significant differences between E198K and E200K are also consistent with the severity difference between the two variants^5, 35^.

**Figure 1:**
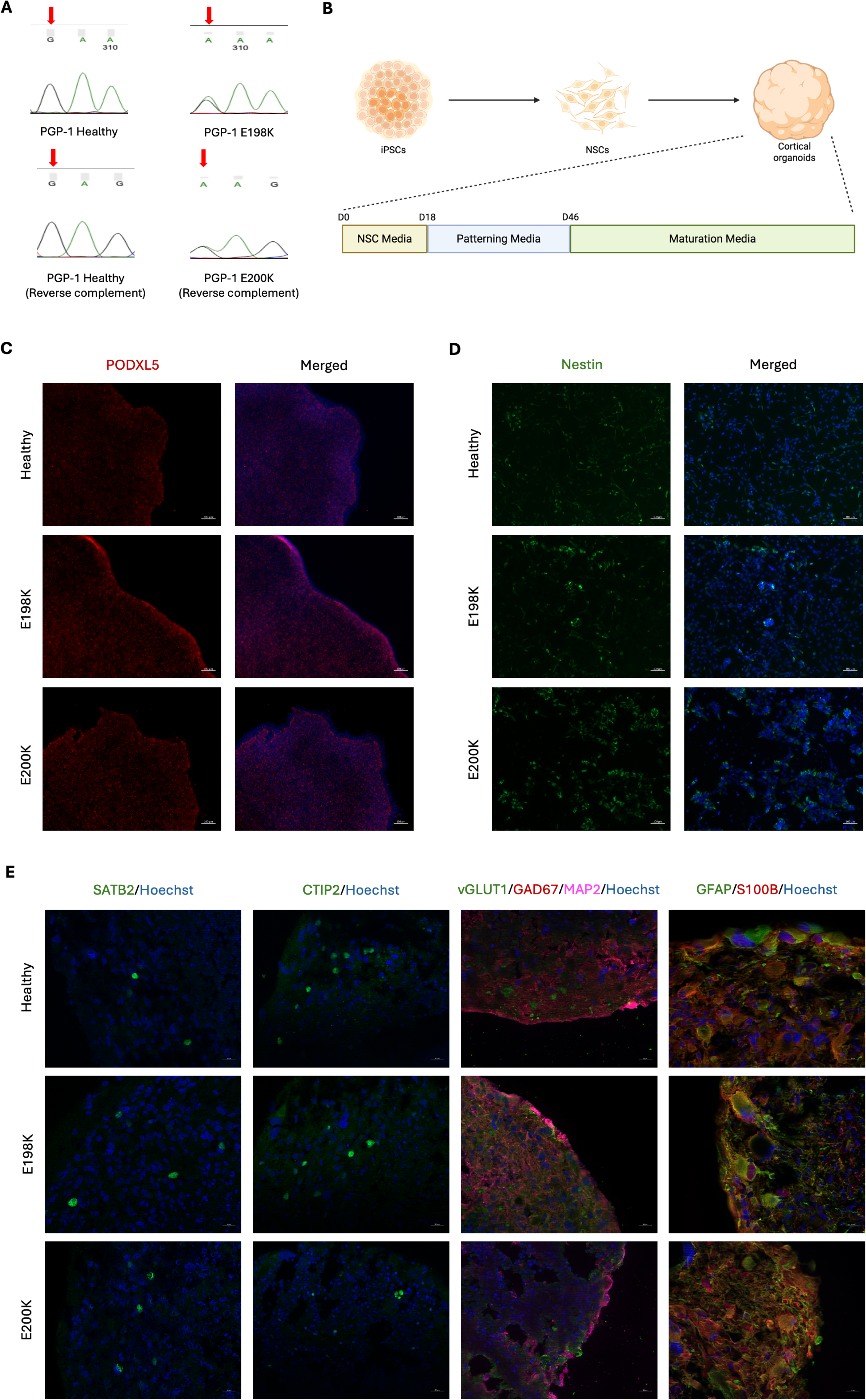
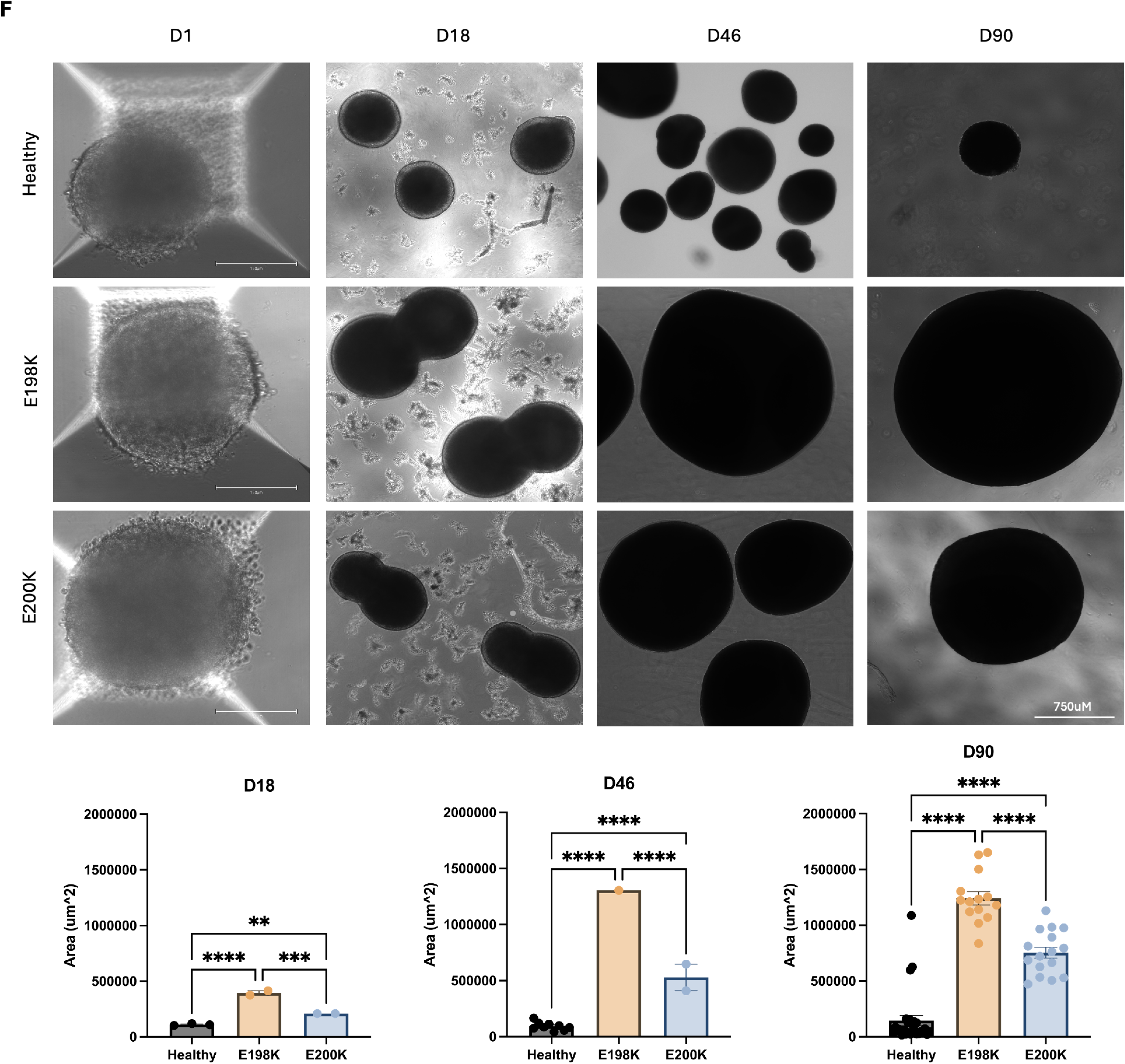
Generation of cerebral cortical-like organoids from iPSCs of PGP-1 allelic series. (A) Sanger sequencing of *PPP2R5D* exon 5 confirms heterozygous guanosine and adenosine peaks in Healthy (left), E198K (right, top) and E200K (right, bottom) lines. (B) Schematic of iPSC differentiation into NSC followed by NSC differentiation into CO. (C) Representative 10X images of Healthy, E198K, and E200K iPSCs positive for PODXL5. Scale bar represents 100um. (D) Representative 10X images of Healthy, E198K, and E200K NSCs positive for Nestin. Scale bar represents 100um. (E) Representative 40X images of Healthy, E198K, and E200K COs positive for SATB2, CTIP2, GAD67, vGLUT1, MAP2, S100B, and GFAP. Scale bar represents 25um (40X). (F) Representative 4X (D18, D46, and D90) and 20X (D1) images of Healthy, E198K, and E200K. Scale bar represents 750um (4X) and 150um (20X). Quantification of CO sizes by ImageJ. One-way ANOVA. * p ≤ 0.05; ** p ≤ 0.01; *** p ≤ 0.001; **** p ≤ 0.0001.

### RNA sequencing results reveal transcriptomic changes in neurodevelopment and transcription factors in mutant organoids

To better understand the role of *PPP2R5D* mutations in human neuronal cells, we performed RNA sequencing on iPSCs, NSCs, early (d18), mid (d46), and late (d85 – d90) stages of COs in all cell lines to examine changes induced by the mutations on a transcriptional level. First, we identified differentially expressed genes (DEGs) between Healthy and the mutants and examined enrichment of neuronal markers in all cell lines across all stages (**Figure 2A**). COs in all cell lines demonstrated upregulation in the excitatory and inhibitory markers when compared to iPSCs and NSCs, and that late-stage COs demonstrated the highest expression of neuronal markers regardless of mutation status. Further, the enrichment analysis of stemness markers (**Supplement Figure 2A**) demonstrated upregulation in iPSCs in all cell lines and the astrocyte markers enrichment analysis (**Supplement Figure 2A**) demonstrated upregulation in late-stage COs in all cell lines. Together, these results showed that the development of COs are consistent with RT-qPCR and ICC results and the model is able to recapitulate different neuronal populations in the cerebral cortex.

**Figure 2:**
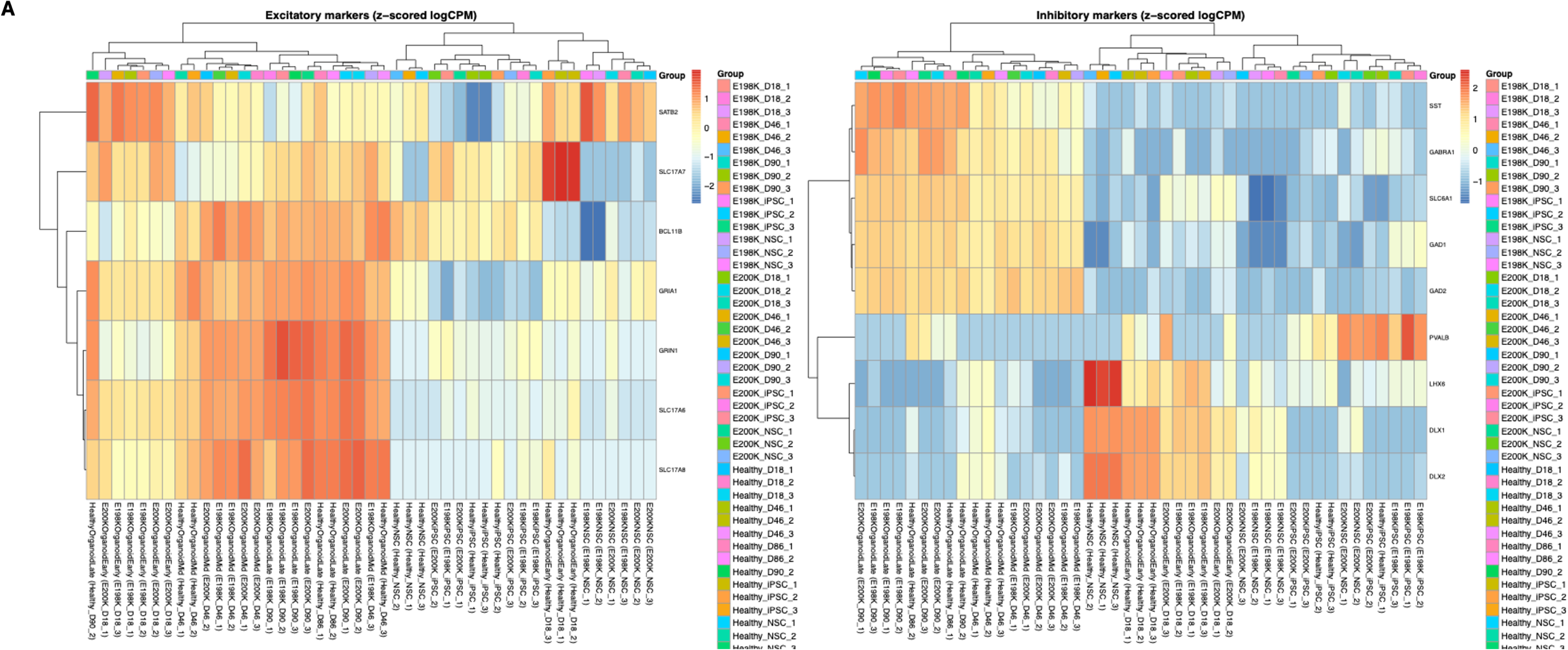
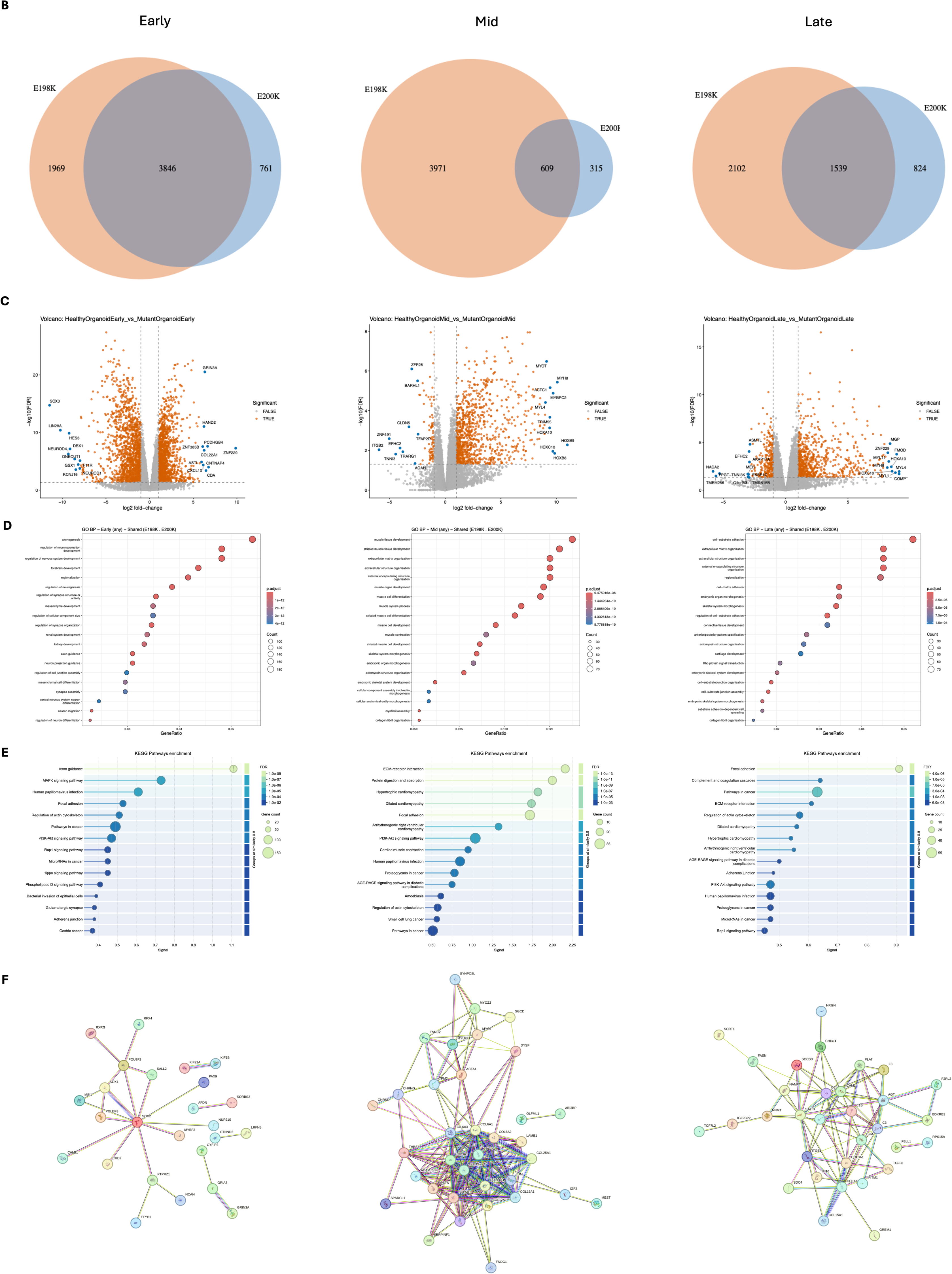
Transcriptomic changes in neurodevelopment and transcription factors are observed in mutant organoids. (A) Excitatory markers enrichment heatmap (left) and inhibitory markers enrichment heatmap (right) of Healthy, E198K, and E200K iPSCs, NSCs, early, mid, and late stages of organoids. (B) Venn diagrams of overlapping dysregulated DEGs between E198K and E200K organoids in early (left), mid (middle), and late (right) stages. (C) Volcano plots of Healthy against mutant organoids in early (left), mid (middle), and late (right) stages. (D) GO term analyses of dysregulated DEGs of Healthy against mutant organoids in early (left), mid (middle), and late (right) stages. (E) KEGG pathway analyses of dysregulated DEGs of Healthy against mutant organoids in early (left), mid (middle), and late (right) stages. (F) STRING analyses of dysregulated DEGs of Healthy against mutant organoids in early (left), mid (middle), and late (right) stages.

In early stage, there are 3846 shared DEGs (58.49%) between the variants, 1969 DEGs unique to E198K, and 761 unique to E200K. In mid stage, there are 609 shared DEGs (12.44%) between the variants, 3971 DEGs unique to E198K, and 315 DEGs unique to E200K. In late stage, there are 1539 shared DEGs (34.47%) between the variants, 2102 DEGs unique to E198K, and 824 DEGs unique to E200K (**Figure 2B**). Volcano plots of early-stage COs showed upregulation of genes associated with neurodevelopment and synaptic function and downregulation of genes associated with neural progenitor development and neuronal lineage specification. Mid-stage COs showed upregulation of *HOX*-regulated body patterning and positional identity and downregulation of cell polarity, cell adhesion, and differentiation states. Late-stage COs showed upregulation, continuing from mid-stage, *HOX*-regulated body posterior mesoderm-derived development, and downregulation of GPCR-mediated signaling and tissue-environment communication (**Figure 2C**; **Table 1**).

**Table 1:**
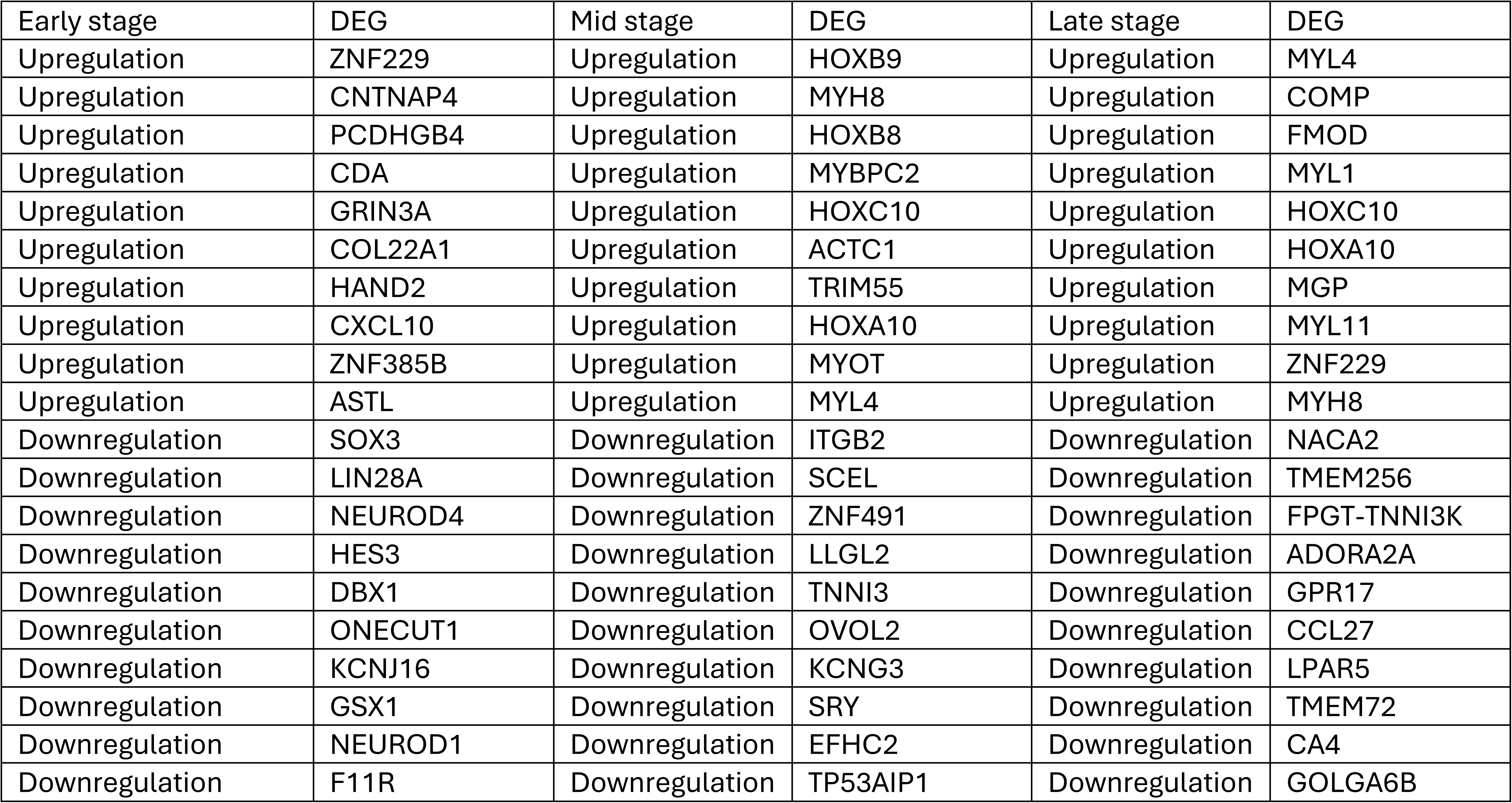
Top 20 DEGs of Healthy and mutant organoids in Early, Mid, and Late stages.

| Early stage | DEG | Mid stage | DEG | Late stage | DEG |
| --- | --- | --- | --- | --- | --- |
| Upregulation | ZNF229 | Upregulation | HOXB9 | Upregulation | MYL4 |
| Upregulation | CNTNAP4 | Upregulation | MYH8 | Upregulation | COMP |
| Upregulation | PCDHGB4 | Upregulation | HOXB8 | Upregulation | FMOD |
| Upregulation | CDA | Upregulation | MYBPC2 | Upregulation | MYL1 |
| Upregulation | GRIN3A | Upregulation | HOXC10 | Upregulation | HOXC10 |
| Upregulation | COL22A1 | Upregulation | ACTC1 | Upregulation | HOXA10 |
| Upregulation | HAND2 | Upregulation | TRIM55 | Upregulation | MGP |
| Upregulation | CXCL10 | Upregulation | HOXA10 | Upregulation | MYL11 |
| Upregulation | ZNF385B | Upregulation | MYOT | Upregulation | ZNF229 |
| Upregulation | ASTL | Upregulation | MYL4 | Upregulation | MYH8 |
| Downregulation | SOX3 | Downregulation | ITGB2 | Downregulation | NACA2 |
| Downregulation | LIN28A | Downregulation | SCEL | Downregulation | TMEM256 |
| Downregulation | NEUROD4 | Downregulation | ZNF491 | Downregulation | FPGT-TNNI3K |
| Downregulation | HES3 | Downregulation | LLGL2 | Downregulation | ADORA2A |
| Downregulation | DBX1 | Downregulation | TNNI3 | Downregulation | GPR17 |
| Downregulation | ONECUT1 | Downregulation | OVOL2 | Downregulation | CCL27 |
| Downregulation | KCNJ16 | Downregulation | KCNG3 | Downregulation | LPAR5 |
| Downregulation | GSX1 | Downregulation | SRY | Downregulation | TMEM72 |
| Downregulation | NEUROD1 | Downregulation | EFHC2 | Downregulation | CA4 |
| Downregulation | F11R | Downregulation | TP53AIP1 | Downregulation | GOLGA6B |

We performed gene ontology analysis (**Figure 2D; Supplement Figure 2C**) and found that DEGs were dysregulated in terms associated with axonogenesis (*GO*: 0007409), regulation of neuron projection development (*GO*:0010975), regulation of nervous system development (*GO*: 0051960), forebrain development (*GO*: 0030900), regionalization (*GO*: 0003002), regulation of neurogenesis (*GO*: 0050767), regulation of synapse structure or activity (*GO*: 0050803), regulation of synapse organization (*GO*: 0050807), axon guidance (*GO*: 0007411), neuron projection guidance (*GO*: 0097485), synapse assembly (*GO*: 0007416), central nervous system neuron differentiation (*GO*: 0021953), neuron migration (*GO*: 0001764), regulation of neuron differentiation (*GO*: 0045664), embryonic organ morphogenesis (*GO*: 0048562), cellular component assembly involved in morphogenesis (*GO*: 0010927), cellular anatomical entity morphogenesis (*GO*: 0032989), cell-substrate adhesion (*GO*: 0031589), extracellular matrix organization (*GO*: 0030198), extracellular structure organization (*GO*: 0043062), external encapsulating structure organization (*GO*: 0045229), cell-matrix adhesion (*GO*: 0007160), regulation of cell-substrate adhesion (*GO*: 0010810), Rho protein signal transduction (*GO*: 0007266), cell-substrate junction organization (*GO*: 0150115), cell substrate junction assembly (*GO*: 0007044), and substrate adhesion-dependent cell spreading (*GO*: 0034446).

To focus on the network of differential genes, we performed a KEGG pathway analyses (**Figure 2E**) and found that DEGs were dysregulated in pathways of axon guidance (hsa: 04360), MAPK signaling pathway (hsa: 04010), focal adhesion (hsa: 04510), pathways in cancer (hsa: 05200), PI3K-Akt signaling pathway (hsa: 04151), Rap1 signaling pathway (hsa: 04015), microRNAs in cancer (hsa: 05206), Hippo signaling pathway (hsa: 04390), phospholipase D signaling pathway (hsa: 04072), glutamatergic synapse (hsa:04724), adherens junction (hsa: 04520), ECM-receptor interaction (hsa: 04512), and complement and coagulation cascades (hsa: 04610).

To assess functional protein associated networks, we performed STRING analyses (**Figure 2F**) and found that DEGs showed strong correlation to the *SOX* family (*SOX1*, *SOX2*) and the *POU3F* family (*POU3F2*, *POU3F3*) and strong clustering to the COL family (*COL1A1*, *COL1A2*, *COL3A1*, *COL5A1*, *COL6A1*, *COL6A2*, *COL6A3*, *COL12A1*, *COL14A1*, *COL16A1*, *COL25A1*.

### Dysfunction of B56δ affects phosphorylation of S6, but not AKT

To better understand previously proposed PP2A-B56δ substrates and how they were affected by the mutations in *PPP2R5D* in neuronal development, we examined phosphorylation states of AKT and S6 ribosomal protein, known downstream targets of PP2A-B56δ in the PI3K/AKT-mTOR signaling pathway^15–17^. First, we examined the expression of B56δ in d86 COs (**Figure 3A**). E198K and E200K are missense mutations that do not cause truncation to the translation of the protein, however, expression of B56δ has a trend of lower expression in E198K with a 0.87 fold change (*p* = 0.3118) and is significantly decreased in E200K with a 0.56 fold-change (p = 0.0038). This result suggests there could be different pathogenic mechanisms underlying the two variants. We then examined the phosphorylation state of AKT at T308 as it is previously reported as a direct substate of PP2A-B56δ in the PI3K/AKT-mTOR signaling pathway^17^ (**Figure 3B**). In contrast to previous publications, Western blot results showed there was no significant changes at AKT T308 between Healthy and E198K with a 0.768 fold-change (*p* = 0.2960), or between Healthy and E200K with a 1.335 fold-change (*p* = 0.4134). However, the significant difference between E198K and E200K suggests different molecular pathological outcome between the two variants. When examining the phosphorylation state of S6 ribosomal protein as the functional downstream target (**Figure 3C**), though, we observed a significant decrease in phosphorylation of S6 at S235: S236 for E198K with a 0.282 fold-change (*p* = 0.0008) but a significant increase for E200K with a 2.158 fold-change (*p* < 0.0001). This is also in contract to previously reported increased phosphorylation state of S6 at this site in E198K in HEK293 cells^14^.

**Figure 3:**
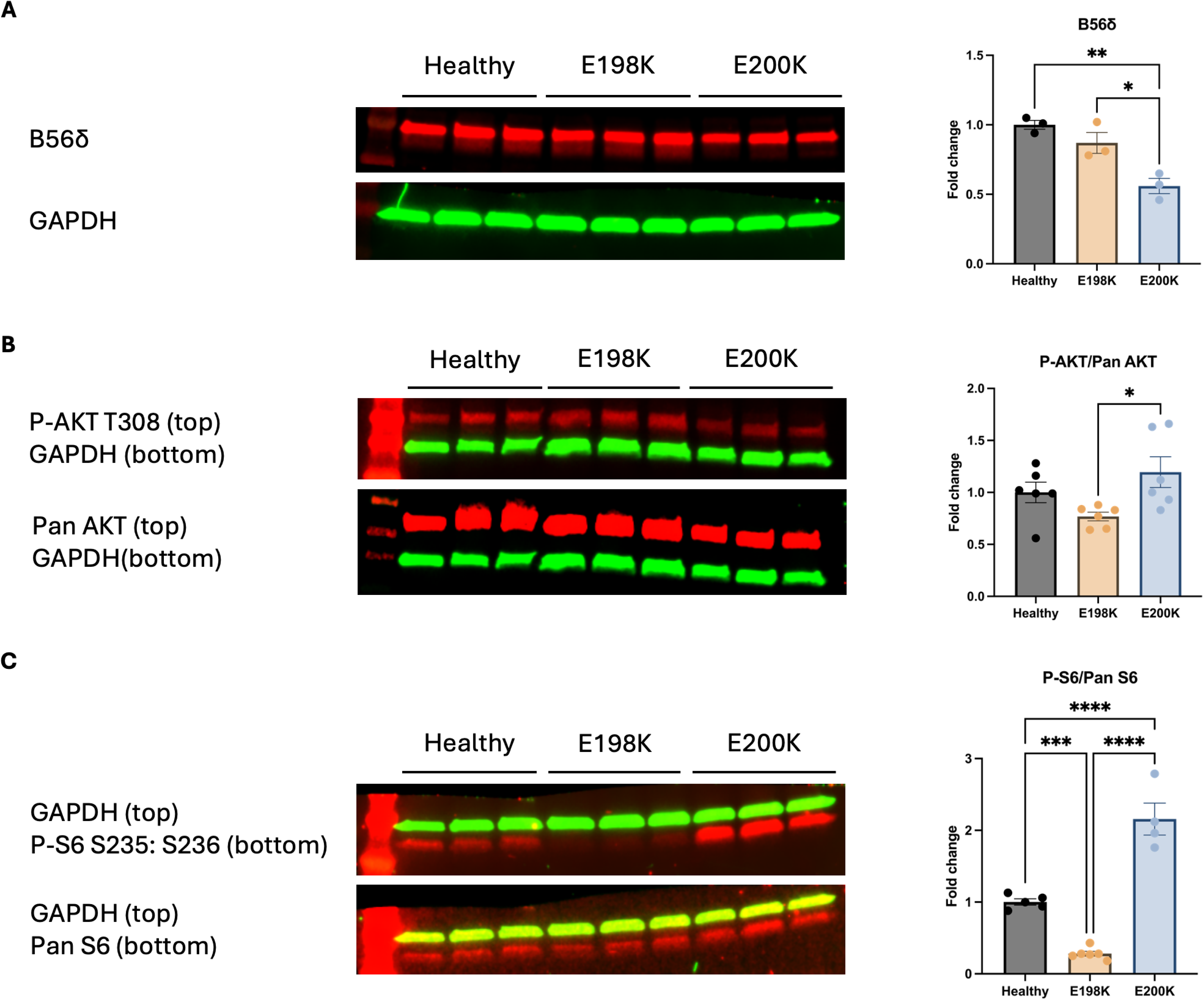
Phosphorylation of downstream targets are affected in mutant organoids. (A) Western Blot of PPP2R5D in Healthy, E198K, and E200K COs with GAPDH housekeeping. Quantification of WB bands by Empiria Studios. One-way ANOVA. * p ≤ 0.05; ** p ≤ 0.01; *** p ≤ 0.001; **** p ≤ 0.0001. (B) Western Blot of Phospho-AKT T308 and Pan AKT in Healthy, E198K, and E200K COs with GAPDH housekeeping. Quantification of WB bands by Empiria Studios. One-way ANOVA. * p ≤ 0.05; ** p ≤ 0.01; *** p ≤ 0.001; **** p ≤ 0.0001. (C) Western Blot of Phospho-S6 S235:S236 and Pan S6 in Healthy, E198K, and E200K Cos with GAPDH housekeeping. Quantification of WB bands by Empiria Studios. One-way ANOVA. * p ≤ 0.05; ** p ≤ 0.01; *** p ≤ 0.001; **** p ≤ 0.0001.

Our group previously reported that TH-positive cells were severely impacted by the mutations in B56δ^4^, suggesting inhibitory neurons could be direct substrates of the subunit. However, when we examined the phosphorylation state of GABA B R2, a previously reported direct substrate of PP2A- B56δ, there was no significant change between healthy and the mutants (**Supplement Figure 3**). Yet, the extremely low expression of Pan GABA B R2 in E200K suggests post-synaptic and GABAergic neurons could be impacted by mutations in *PPP2R5D*.

In addition, to identify potential downstream substrates and to better understand the differences in function between the two mutated phosphatases, we performed phospho-kinase arrays on d123 COs (**Supplement Figure 6B - C**). Among 39 targets examined, 33 targets demonstrated decreases in their phosphorylation state in the mutants.

### Longitudinal network analyses suggest network dysregulation in mutant organoids

To investigate how mutations in *PPP2R5D* affect neuronal function and network activity, we used high-density microelectrode array (HD-MEA; Maxwell Biosystems) to measure the electrophysiological phenotypes of COs across multiple timepoints on chip. Neuronal unit recordings were spike-sorted to identify putative neurons, and the number of detected units was tracked across recording days (**Figure 4A**). The lower unit count at day-on-chip (DOC) 7 reflects an initial stabilization period, after which COs exhibited consistent and comparable activity across conditions (**Supplement Figure 4**), ensuring reliable cross-condition comparisons. DOC 14 was selected as an optimal timepoint for downstream analysis. Representative spike-averaged waveform templates from the top 10% amplitude putative neurons ranked by amplitude confirmed biologically valid action potential shapes, reflecting the dominant ionic currents at the extremum channel of each electrode (**Figure 4B**). Functional amplitude maps revealed the spatial distribution of neuronal activity across the chip, with the locations of identified putative neurons overlaid, highlighting regions of concentrated neuronal hotspots (**Figure 4C**). Raster plots of coordinated network activity showed that healthy COs display well-regulated bursting with a higher network burst rate, while E198K COs exhibited greater neuronal recruitment and larger burst amplitudes, followed by E200K COs, suggesting a mutation-dependent increase in network excitability (**Figure 4D**). Quantification of unit-level metrics revealed that E200K COs exhibited a significantly higher mean firing rate compared to Healthy COs (**Figure 4E-i**, *p* = 0.00219), while mean spike amplitude remained comparable across all three conditions (**Figure 4E-ii**). At the network burst level, E198K COs showed a significantly lower network burst rate compared to both Healthy and E200K COs (**Figure 4F-i**, *p* = 0.02242 and *p* = 0.01173, respectively). Despite this reduced burst rate, E198K COs displayed significantly longer burst durations compared to both Healthy and E200K COs (**Figure 4F-ii**, *p* = 0.00042 and *p* = 0.001498, respectively), as well as a greater number of spikes per burst (**Figure 4F-iii**, *p* = 0.003271 and *p* = 0.001187, respectively), indicative of larger and more prolonged bursting events consistent with the more severe E198K phenotype. E200K COs showed intermediate burst durations and spikes per burst, both significantly different from Healthy COs (**Figure 4F-ii**, *p* = 0.003865; **Figure 4F-iii**, *p* = 0.02897). Waveform analysis revealed that peak-to-valley time was significantly altered in E198K COs compared to Healthy controls (**Figure 4G-i**, *p* = 0.04431), while repolarization slope was significantly different in E200K COs compared to Healthy controls (**Figure 4G-ii**, *p* = 0.0238), suggesting mutation-specific changes in action potential morphology, potentially reflecting altered phosphoregulation of ion channels downstream of disrupted PP2A-B56δ activity. Collectively, these findings reveal distinct and mutation-specific electrophysiological phenotypes in *PPP2R5D*-mutant COs, both mutant lines exhibit dysregulated network dynamics, characterized by less frequent but larger and more prolonged bursting events, suggestive of impaired network regulation.

**Figure 4:**
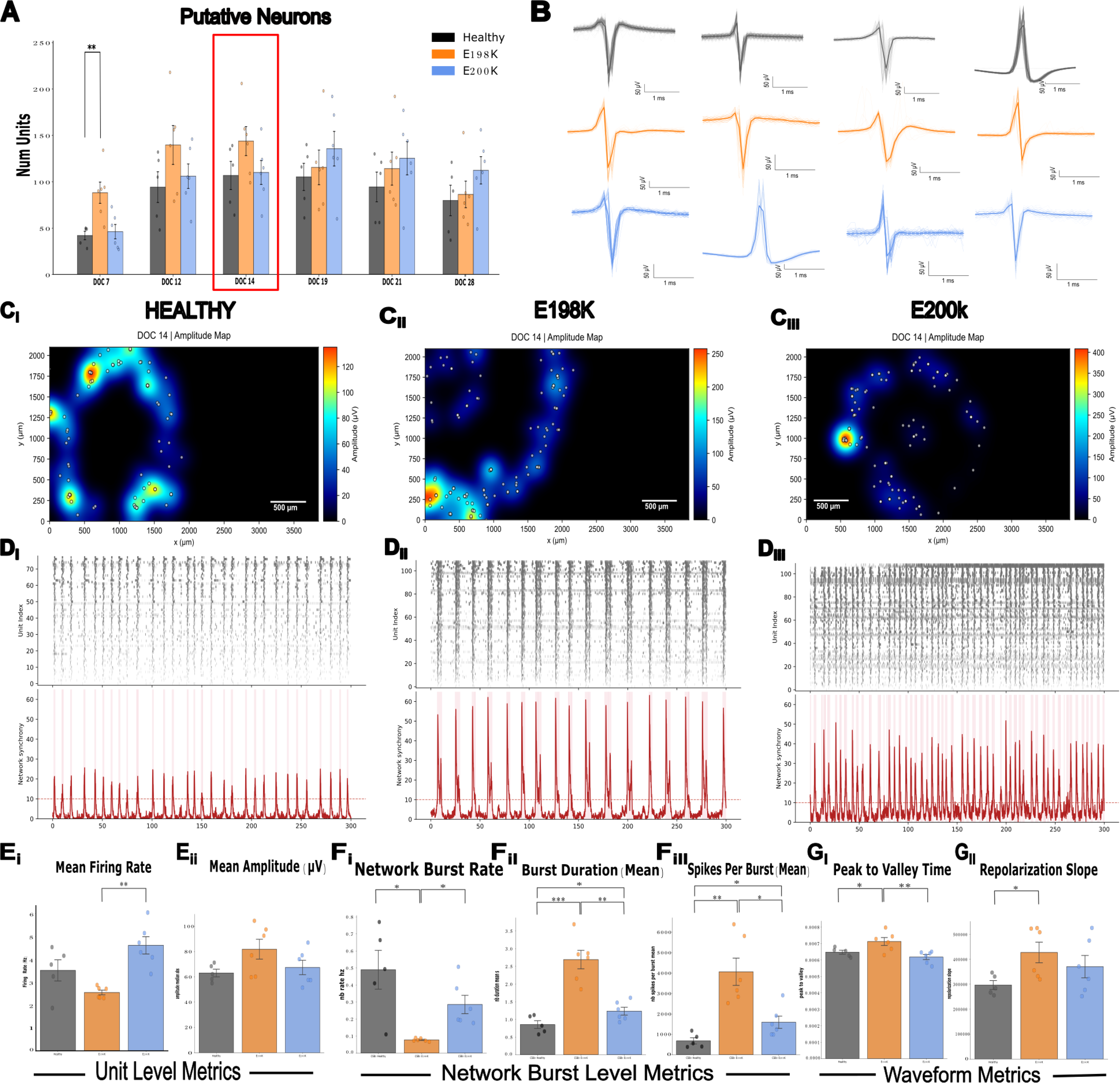
Mutant CO variants exhibit distinct electrophysiological phenotypes compared to Healthy CO controls. (A) Quantification of putative neurons detected across developmental time points (DOC 7-28) in Healthy, E198K, and E200K cultures. DOC 14 (red box) was selected for downstream analyses. Data are shown as mean ± SEM. (B) Representative spike-averaged waveform templates from the top 10% of putative neurons ranked by spike amplitude at DOC 14. Waveforms display canonical action potential morphologies recorded at the extremum channel of an identified putative neuron. Scale bars: 50 µV, 1 ms. (C) Spatial spike amplitude maps at DOC 14 for (Ci) Healthy, (Cii) E198K, and (Ciii) E200K cultures. Color scale represents spike amplitude (µV); white dots indicate location of the putative neurons. Scale bar: 500 µm. (D) Representative raster plots (top) and corresponding network synchrony traces (bottom) at DOC 14 for (Di) Healthy, (Dii) E198K, and (Diii) E200K cultures. (E) Quantification of electrophysiological metrics at DOC 14. Unit level: (Ei) Mean firing rate. (Eii) Mean spike amplitude. Data are presented as mean ± SEM with individual replicates overlaid. Statistical significance is indicated as * p < 0.05, ** p < 0.01, *** p < 0.001. (F) Quantification of electrophysiological metrics at DOC 14. Network burst level: (Fi) Network burst rate. (Fii) Burst duration. (Fiii) Spikes per burst (mean). Data are presented as mean ± SEM with individual replicates overlaid. Statistical significance is indicated as * p < 0.05, ** p < 0.01, *** p < 0.001. (G) Quantification of electrophysiological metrics at DOC 14. Waveform metric: (Gi) Peak-to-valley time. (Gii) Repolarization slope. Data are presented as mean ± SEM with individual replicates overlaid. Statistical significance is indicated as * p < 0.05, ** p < 0.01, *** p < 0.001.

### Rapamycin improves AKT/mTOR-downstream and electrophysiological phenotypes of mutant organoids

Previous studies have shown that Rapamycin is an effective inhibitor in AKT-mTOR signaling pathway in neurons^25, 26^. To investigate whether Rapamycin could rescue or restore JS-related phenotypes, we tested Rapamycin on d123 COs. COs received Rapamycin at 20 nM/organoid and were incubated at 37°C between 45 minutes and 1 hour to allow full take up of the drug without compromising the decay of the molecule.

We first examined the phosphorylation of S6 at S235:S236 as it demonstrated significant changes in untreated COs. Quantification of Western blot results showed reproducible phenotypes in the DMSO-treated COs where there is a significant decrease in E198K (*p* = 0.0211) and increase in E200K (*p* = 0.1170). (**Figure 5A**). Rapamycin-treated COs demonstrated a decreasing trend of S6 phosphorylation in all cell lines. Healthy Rapamycin-treated COs have a 0.537 fold-change compared to DMSO-treated COs, E198K Rapamycin-treated COs have a 0.670 fold-change compared to DMSO-treated COs, and E200K Rapamycin-treated COs have a 0.423 fold-change compared to DMSO-treated COs. E198K Rapamycin-treated COs have a 0.15 fold-change (*p* = 0.01), and E200K Rapamycin-treated COs have a 0.67 fold-change (*p* = 0.8035) compared to Healthy DMSO-treated COs. Overall, Rapamycin treatment resulted in decreased phosphorylation of S6 at S235:S236 in all cell lines, as expected, but it resulted in an inverse rescue effect in E198K and E200K.

**Figure 5:**
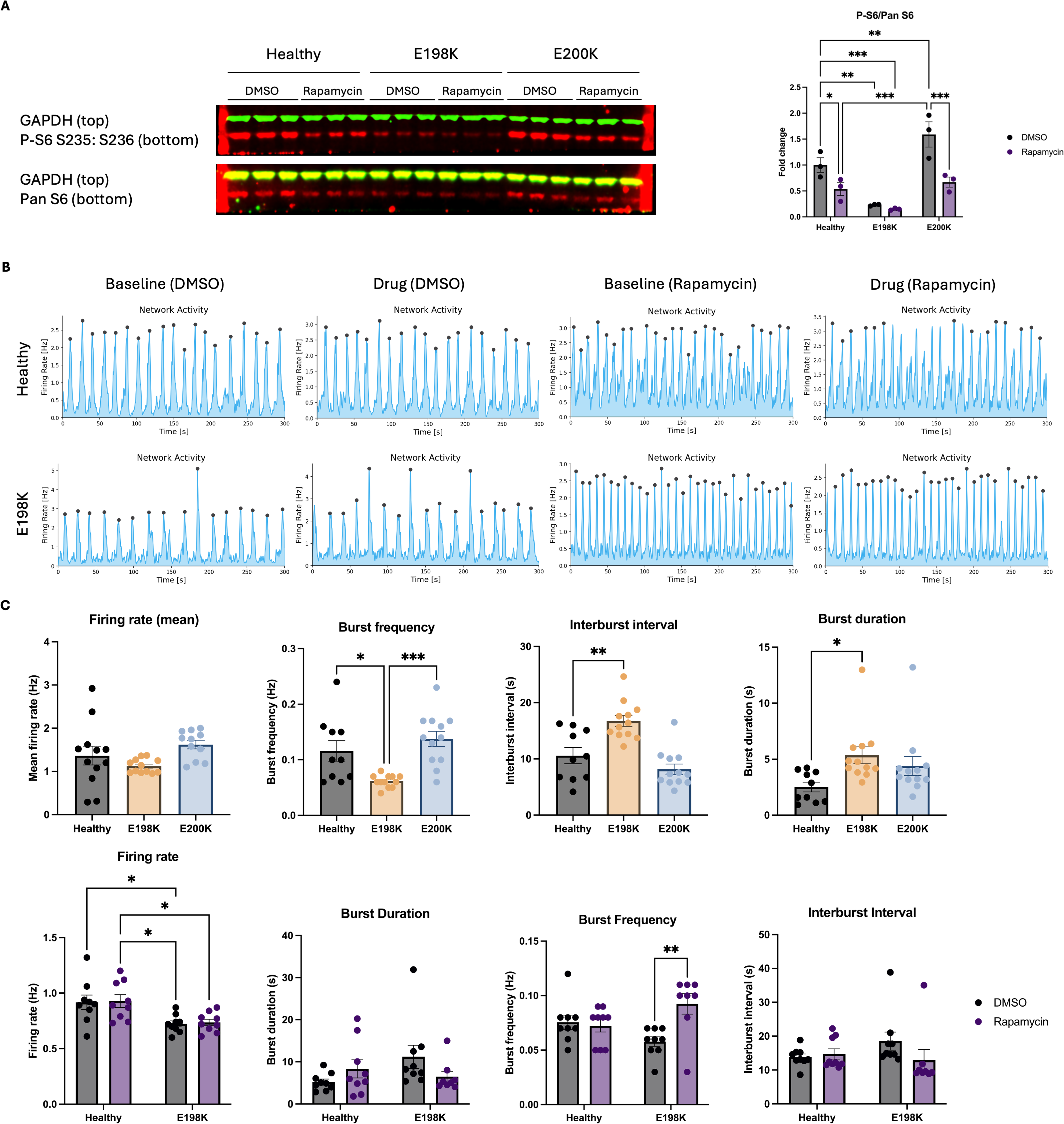
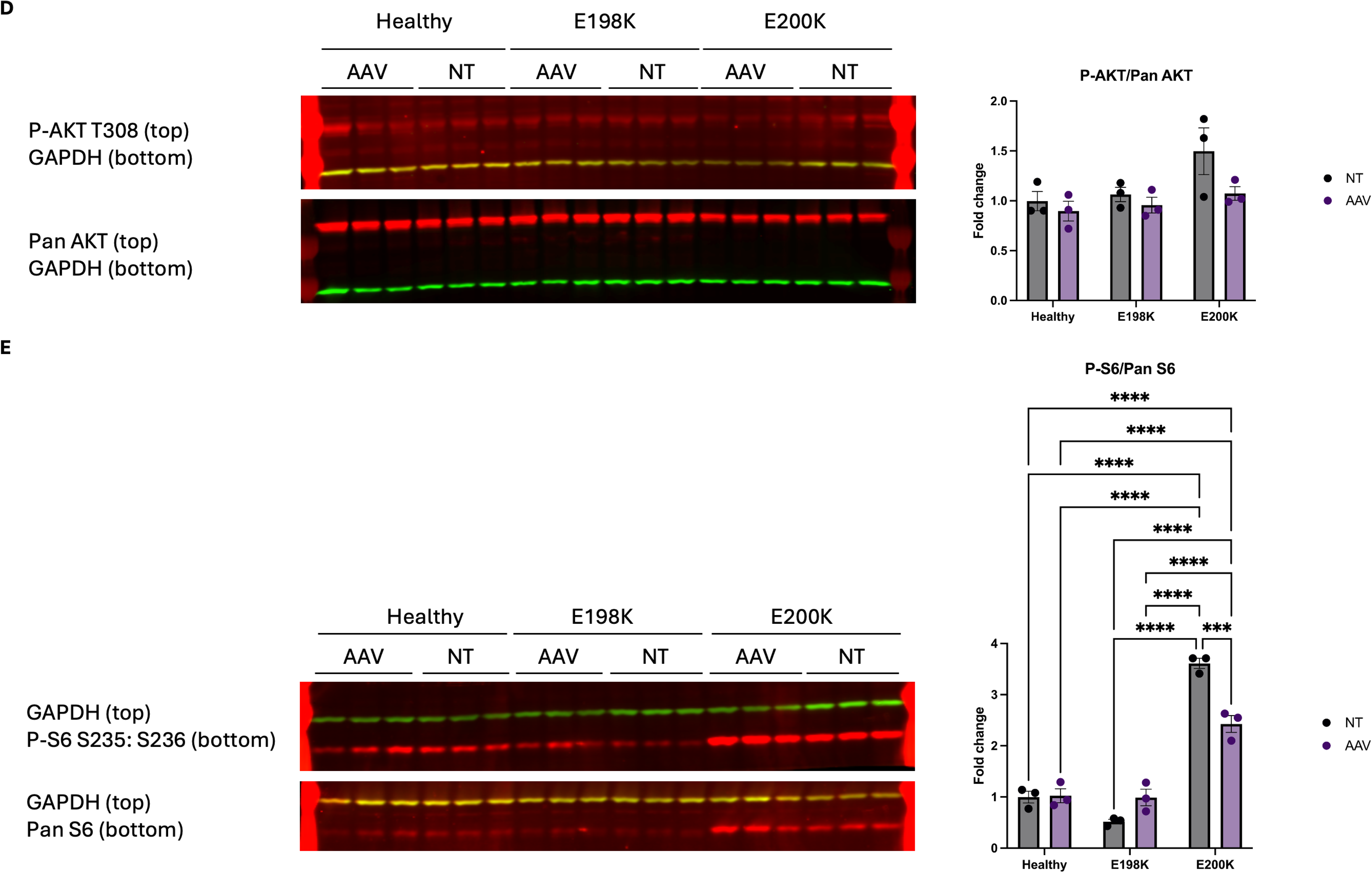
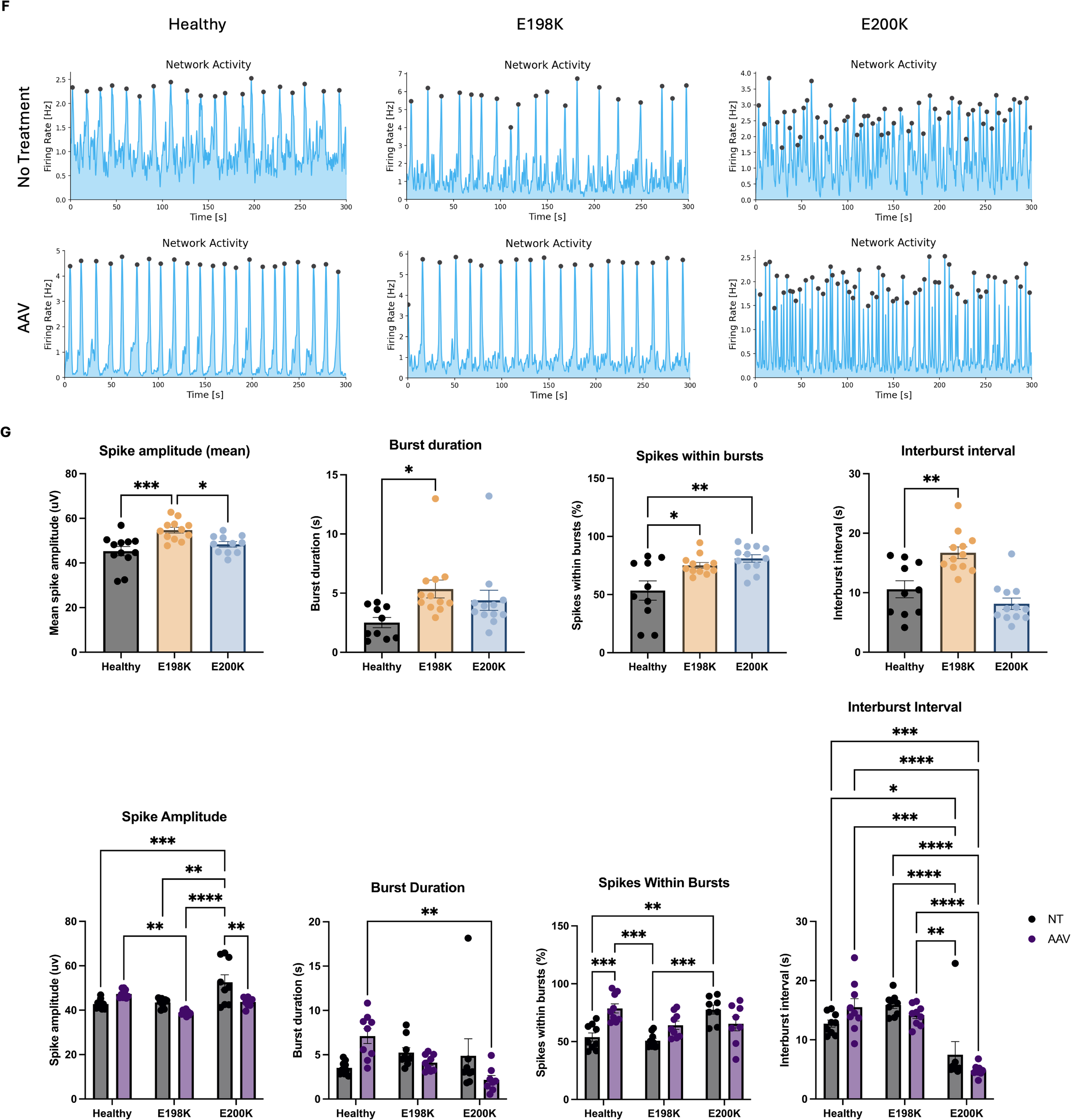
Pharmaceutical and genetic intervention rescued different phenotypes in the variants. (A) Western Blot of Phospho-S6 S235:S236 and Pan S6 in Healthy, E198K, and E200K COs with GAPDH housekeeping. Quantification of WB bands by Empiria Studios. Two-way ANOVA. * p ≤ 0.05; ** p ≤ 0.01; *** p ≤ 0.001; **** p ≤ 0.0001. (B) Representative images of Network Activity plot of Healthy, E198K, and E200K COs on Maxwell MaxTwo MEA system. (C) Quantification of electrophysiological metrics at DOC 14, 16, and 19. Data are presented as mean ± SEM with individual replicates overlaid. Statistical significance is indicated as * p < 0.05, ** p < 0.01, *** p < 0.001. (D) Western Blot of Phospho-AKT T308 and Pan AKT in Healthy, E198K, and E200K Cos 17 days post transduction with GAPDH housekeeping. Quantification of WB bands by Empiria Studios. One-way ANOVA. * p ≤ 0.05; ** p ≤ 0.01; *** p ≤ 0.001; **** p ≤ 0.0001. (E) Western Blot of Phospho-S6 S235:S236 and Pan S6 in Healthy, E198K, and E200K Cos 17 days post transduction with GAPDH housekeeping. Quantification of WB bands by Empiria Studios. One-way ANOVA. * p ≤ 0.05; ** p ≤ 0.01; *** p ≤ 0.001; **** p ≤ 0.0001 (F) Representative images of Network Activity plot of Healthy, E198K, and E200K COs 37 days post transduction on Maxwell MaxTwo MEA system. (G) Quantification of electrophysiological metrics at DOC 14, 19, and 21. Data are presented as mean ± SEM with individual replicates overlaid. Statistical significance is indicated as * p < 0.05, ** p < 0.01, *** p < 0.001.

To investigate whether Rapamycin can rescue neuronal function, we then used HD-MEA to assess the electrophysiological changes in COs treated with Rapamycin. Because E198K variant displays the most severe symptoms in JS^27^ as well as the most significant changes in HD-MEA assessment, we only used this mutant for Rapamycin treatment. DOC14, 16, and 19 were used in the quantification of HE-MEA parameters. Network activity plots showed no clear changes between firing patterns of baseline and treated COs in all cell lines (**Figure 5B**). Interestingly, Rapamycin increased burst frequency and decreased burst duration and interburst interval in E198K COs (**Figure 5C**). Of these parameters, E198K Rapamycin-treated COs showed burst duration of 6.494s, which is comparable to Healthy DMSO-treated COs burst duration of 5.219s (*p* = 0.9672). E198K Rapamycin-treated COs also displayed interburst intervals of 12.87s, which is comparable to Healthy DMSO-treated COs interburst intervals of 13.82s (*p* = 0.9902). Even though burst frequency in E198K Rapamycin-treated COs surpassed the Healthy DMSO-treated COs, this parameter is still restored in the correct direction to be comparable to the Healthy COs.

### Overexpression of PPP2R5D revealed gene dosage sensitivities between the variants

While JS is caused by heterozygous mutation in *PPP2R5D*, the specific pathogenetic etiology remains unclear. To investigate if the mutations in *PPP2R5D* are truly dominant negative, we delivered a healthy, codon-optimized copy of *PPP2R5D* in the mature COs via AAV9 transduction and examined phenotypic changes. A codon-optimized *PPP2R5D-IRES-GFP* was cloned into an AAV backbone under the CMV promoter (**Supplement Figure 5A**). *PPP2R5D* overexpression was validated in *in vitro* and confirmed to increase total *PPP2R5D* expression in HEK293 cells (**Supplement figure 5B, C**). 10^11 total vector genomes (vg) were added directly to the cell culture media of d70 COs. The AAV was confirmed to be delivered to the cells 17 days post-transfection with positive GFP signal (**Supplement figure 5D**). Transcriptomic expression of codon-optimized *PPP2R5D* was confirmed through RT-qPCR in all cell lines (**Supplement Figure 5E**). As expected, protein expression of B56δ showed an increase in Healthy with a 1.71 fold-change (*p* = 0.0003), and in E200K with a 1.587 fold-change (*p* = 0.0080). However, there was no significant increase in B56δ expression in E198K with a 1.187 fold-change (*p* = 0.2843) (**Supplement Figure 5F**). We then examined the phosphorylation state of AKT at T308 in the transduced COs (AAV) (**Figure 5D**). Even though E200K AAV COs displayed a decreasing trend, statistically the phosphorylation levels remained non-significant in both untreated and treated COs, once again suggesting that this site on AKT is not a direct substrate of PP2A- B56δ in the neuronal cell population. However, when examining the phosphorylation state of S6 at S235:S236 in the treated COs (**Figure 5E**), unexpectedly, even though E198K AAV COs did not express significant amount of extra B56δ, the phosphorylation level increased from 0.52 fold-change to 0.99 fold-change in the treated COs and it is now comparable to untreated (NT) Healthy COs. In E200K AAV COs, on the other hand, the phosphorylation level decreased significantly from 3.61 fold-change to 2.43 fold-change.

We then assessed network activity using HD-MEA to assess the COs on the longitudinal network analyses. Network activity plots showed no clear changes between firing patterns of treated and untreated COs in all cell lines (**Figure 5F)**. Quantification of activity scans and network analyses suggest different rescued parameters in the variants (**Figure 5G**). Both E198K and E200K AAV COs showed same trends in spike amplitude, burst duration, and interburst interval. For E198K AAV COs, even though the changes in parameters were in the correct direction to be comparable to Healthy NT COs, none of the parameters was significantly rescued. On the other hand, E200K AAV COs fired with an average of 43.73 μ V spike amplitude and 65.46% spikes within burst, which are comparable to Healthy NT COs at 42.79 μ V spike amplitude (*p* >0.9999) and 53.90% spikes within burst (*p* = 0.5220).

## Discussion

The growing and aging patient population of JS speaks to an urgent and critical need to understand and characterize the disease. However, the lack of a human neuronal model has been a major challenge in the investigation of the *PPP2R5D*-induced phenotype and mechanisms. Here, we develop a human 3D neuronal model to characterize the molecular and cellular phenotypes of JS. We used COs differentiated from iPSCs of an allelic series and thus provide a more complex model in identifying disease phenotypes in different variants.

Throughout CO development, we observed morphological, transcriptomic, protein function, and electrophysiological phenotypes in the mutants. Morphologically, the mutant COs demonstrated significantly increased sizes compared to the Healthy COs and this observation is consistent with the clinical symptoms of macrocephaly and frontal bossing in JS patients. On the transcriptomic level, the mutant COs’ top DEGs demonstrated the shared themes of neurodevelopment, transcription factor, and protein activities. Zinc finger (*ZNF385B*, *ZNF229*, *ZFP28*, *ZNF491*), the *HOX* family (*HOXA10*, *HOXC10*, *HOXB9*, *HOXB8*), the *SOX* family (*SOX1*, *SOX2*), *POU3F* family (*POU3F2*, *POU3F3*), and the *COL* family (*COL1A1*, *COL1A2*, *COL3A1*, *COL5A1*, *COL6A1*, *COL6A2*, *COL6A3*, *COL12A1*, *COL14A1*, *COL16A1*, *COL25A1*) are the most common transcription factors affected. This suggests that *PPP2R5D* could be involved in regulation of transcription, similar to previously reported interaction and involvement of PP2A A subunits with IntS8 and RNA Polymerase II recruitment^29^. In KEGG pathway analyses, PI3K/AKT-mTOR signaling pathway showed to be dysregulated in all three stages, validating the direct impact of *PPP2R5D* mutations on this pathway as previously reported. In addition, MAPK, Rap1, Hippo, and Phospholipase D signaling pathways were also identified to be dysregulated in the mutants, providing potential pathways that are directly impacted by the mutations. Late-stage organoid STRING analysis showed no clustering of specific gene family, but the variety of genes in the network could be a result of affected transcription factors in the early development, further enhancing that JS is a neurodevelopmental disorder. Furthermore, there are overall more DEGs unique to E198K than to E200K across all stages. This suggests a more global change in E198K and is consistent with the different severity observed clinically between the two variants. On the protein function level, the mutant COs demonstrated significant differences between each other and are inconsistent with previous studies^14^ on phosphorylation of S6 ribosomal protein. One possible explanation is the different mitotic state of the cells that at the same stage of development, E198K is in a more quiescent state and E200K is in a more active state. This difference also speaks the need to use cell-type specific models for studying disease. In addition, while S6 indicates changes in the downstream outcome of AKT-mTOR signaling pathway, the non-significant phosphorylation states of AKT at T308 suggest that the dysfunction of B56δ does not directly impair the substrate specificity of AKT. This implies that there is another potential substrate that is being impaired along this signaling pathway. Finally, electrophysiologically, longitudinal network analysis of Cos revealed dysregulated network activity in E198K and E200K, with both mutants demonstrating distinct firing patterns compared to Healthy COs. E198K COs were characterized by the largest and the most prolonged bursting events among the three cell lines, while E200K COs were characterized by the most frequent firing. Although the specific neuronal firing networks were different between the two variants and that the underlying cause of this network dysregulation remains unclear, it is highly possible that this is a consequence of E/I imbalance.

Unfortunately, currently there is no cure for JS. However, Rapamycin, as an inhibitor in mTOR signaling pathway, is a strong candidate in many neurological disorder treatments. In our test on the JS model, it demonstrated potential in restoring certain molecular and cellular phenotypes. In the network activity analyses on COs treated with Rapamycin, the small-molecule drug was able to restore firing rate, burst duration, and interburst interval in E198K to be comparable to Healthy DMSO treated COs. However, because the quantification of other parameters suggests no changes, this difference could only be a result of Rapamycin but not specific to JS phenotype. Furthermore, different dosages could induce different restoration levels, and more studies should be conducted in investigating Rapamycin usage in treating JS patients.

Since the discovery of the disease in 2014, JS has been hypothesized to be a dominant negative disorder. However, the pathogenetic mechanisms remain unclear and the question of why different variants display severities across a wide range is still unanswered. Gene complementation of healthy *PPP2R5D* through AAV transduction sheds light on the answers to these questions by suggesting different pathogenic phenotypes between the variants . Following transduction of the AAV-encoding PPP2R5D, protein expression in E198K COs remains statistically unchanged. It was previously reported that PP2A directly interacts with RNA polymerase II and mediate transcription termination^29^ and this could be a possible explanation of the lack of PPP2R5D protein expression. In addition, it was reported that E3 ubiquitin selectively degrades B’ β subunit of PP2A^33^ and this observation could be a proteasome-driven self-regulation of *PPP2R5D*. However, even without significantly increased protein expression in E198K, Western blot results show rescue of phosphorylation of S6 at S235:236. The phospho-kinase array results also show that there are still restored phosphorylation level in Chk-2, p53, p70 S6 kinase, PYK2, RSK, STAT3, and HSP60. Longitudinal network analyses show no clear rescue in E198K, but in E200K in spike amplitude, burst duration, and spikes within bursts. While current evidence is not conclusive in determining the specific pathogenic mechanisms underlying the two variants, it is highly possible that they have different genetic etiology, and further studies should be conducted in providing more sufficient information.

Here, we demonstrated that our human 3D neuronal model was able to recapitulate molecular, cellular, and genetic severity of phenotypes of JS. This study identifies pathways that are significantly manifested, characterized neuronal firing network in different variants of *PPP2R5D* mutation, and tested both cell and gene therapeutic and small-molecule drug approaches.

## Acknowledgments

Support for this project was provided by Jordan’s Guardian Angels, State of California 2018-2019 Budget; SB 840, State of California 2021-2022 Budget; SB 129 #44, NIH eMCB T32, graphics created in biorender.

## Supplemental tables

**Table S1:**
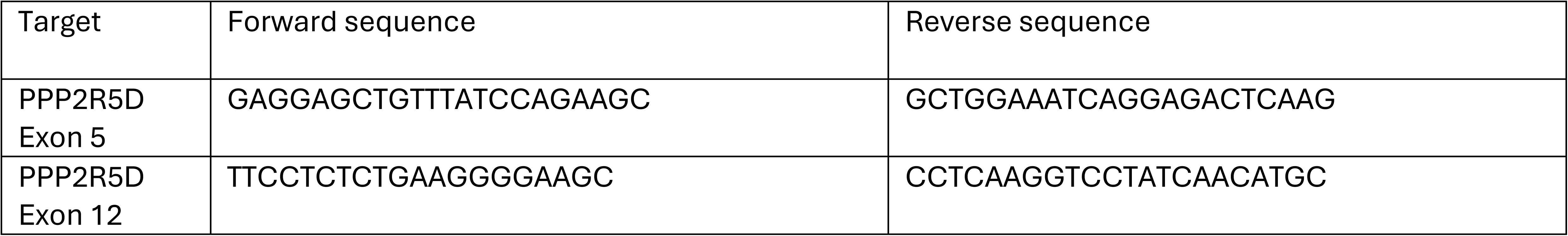
Sanger sequencing primers.

**Table S2:**
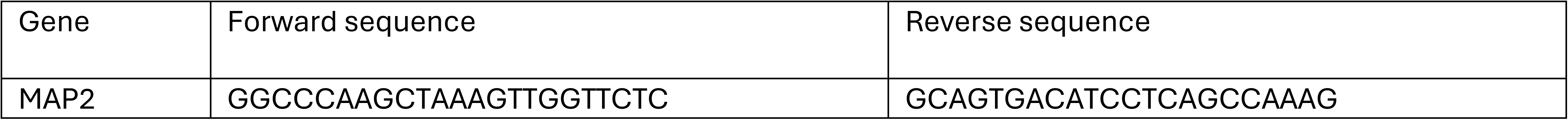
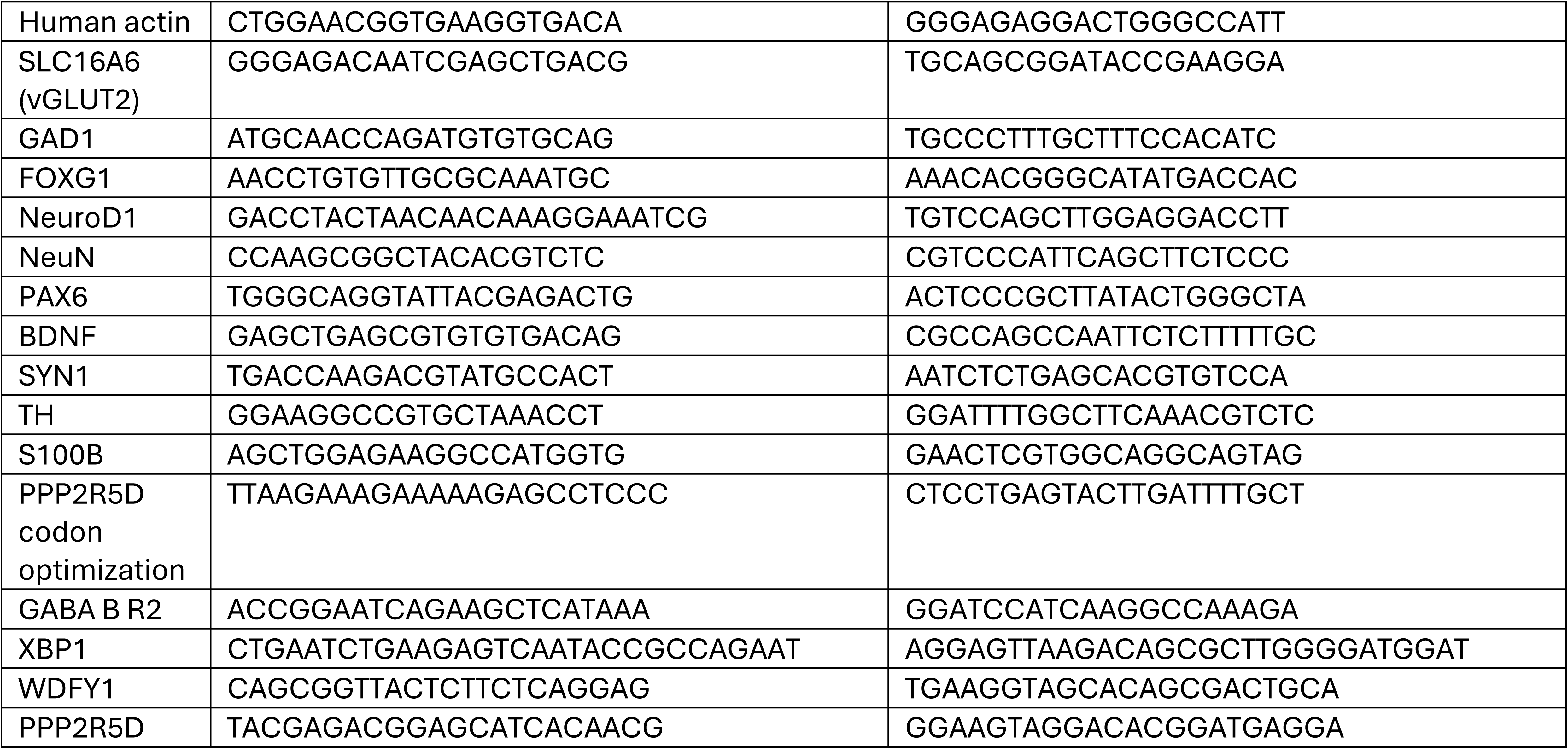
qPCR primers.

**Table S3:** Primary antibodies used in immunocytochemistry staining.

| Primary Antibody | Host | Manufacturer | Catalog Number | Dilution |
| --- | --- | --- | --- | --- |
| PODXL | Mouse | R&D Systems | SC025 | 1:100 |
| Nestin | Rabbit | Abcam | 176571 | 1:100 |
| SATB2 | Mouse | Abcam | AB51502 | 1:50 |
| TBR2 | Rabbit | Millipore | AB2283 | 1:100 |
| CTIP2 | Rat | Abcam | AB18465 | 1:00 |
| TUJ1 | Mouse | Abcam | AB78078 | 1:500 |
| MAP2 | Rabbit | Abcam | AB254264 | 1:1000 |
| NeuN | Guinea pig | Millipore | ABN90 | 1:500 |
| PAX6 | Rabbit | BioLegend | 901301 | 1:400 |
| TH | Chicken | Abcam | AB76442 | 1:250 |
| vGLUT1 | Guinea pig | Millipore | AB5905 | 1:250 |
| S100B | Rabbit | Abcam | AB52642 | 1:100 |
| SOX2 | Rabbit | Cell Signaling | 35795 | 1:400 |
| Nestin | Mouse | Abcam | AB22035 | 1:100 |
| GAD67 | Mouse | Millipore | MAB5406 | 1:100 |
| Nkx2.1 | Rabbit | Abcam | AB76013 | 1:250 |
| GFAP | Mouse | Abcam | AB4648 | 1:100 |
| MAP2 | Chicken | Abcam | AB5392 | 1:100 |

**Table S4:** Secondary antibodies used in immunocytochemistry staining.

| Secondary Antibody | Host | Manufacturer | Catalog Number | Dilution |
| --- | --- | --- | --- | --- |
| Alexa Fluor 594, anti-Rabbit IgG (H+L) | Goat | Thermofisher | A11012 | 1:250 |
| Alexa Fluor 488, anti-Mouse IgG (H+L) | Goat | Thermofisher | A11001 | 1:250 |
| Alexa Fluor 647, anti-Rat IgG (H+L) | Goat | Thermofisher | A21472 | 1:250 |
| Alexa Fluor 647, anti-Guinea Pig IgG (H+L) | Goat | Thermofisher | A21450 | 1:250 |
| Alexa Fluor 488, anti-Rabbit IgG (H+L) | Goat | Thermofisher | A11008 | 1:250 |
| Alexa Fluor 647, anti-Chicken IgG (H+L) | Goat | Thermofisher | A21449 | 1:250 |
| Alexa Fluor 594, anti-Guinea Pig IgG (H+L) | Goat | Thermofisher | A11012 | 1:250 |
| Alexa Fluor 488, anti-Guinea Pig IgG (H+L) | Goat | Thermofisher | A11073 | 1:250 |
| Alexa Fluor 594, anti-Mouse IgG (H+L) | Goat | Thermofisher | A11005 | 1:250 |

**Table S5:** Primary antibodies used in Western blot.

| Primary Antibody | Host | Manufacturer | Catalog Number | Dilution |
| --- | --- | --- | --- | --- |
| Akt (pan) | Rabbit | Cell Signaling | 4691 | 1:1000 |
| Phospho-Akt (Thr308) | Rabbit | Cell Signaling | 13038 | 1:500 |
| S6 Ribosomal protein | Rabbit | Cell Signaling | 2217 | 1:500 |
| Phospho-S6 Ribosomal protein (Ser235/236) | Rabbit | Cell Signaling | 4858 | 1:500 |
| Phospho-GABA B R2 (Ser783) | Rabbit | Abcam | Ab72447 | 1:500 |
| GABA B R2 | Rabbit | Abcam | Ab75838 | 1:500 |
| PPP2R5D | Rabbit | Abcam | Ab188323 | 1:1000 |

**Table S6:** Secondary antibodies used in Western blot.

| Secondary Antibody | Host | Manufacturer | Catalog Number | Dilution |
| --- | --- | --- | --- | --- |
| 800 RD Anti-Mouse | Donkey | Li-Cor | 926-32212 | 1:2000 |
| 680 RD Anti-Rabbit | Donkey | Li-Cor | 926-68071 | 1:2000 |

## Methods

### iPSC differentiation to NSC

Human iPSCs were cultured in feeder-free conditions and differentiated into NSCs through a 12-day process. On day 1, iPSCs were plated in AggreWell 800 plate (STEMCELL Technologies #34811) at the density 3×10^6^ cells per well on a 24-well plate in Embryoid Body (EB) media containing Knockout DMEM/F12 (Gibco #12660012), 15% knockout serum replacer (Gibco #10828010), 100x MEM non-essential amino acids (NEAA) (Gibco #11140050), 100x Glutamax L-glutamine (Glutamax) (Gibco #35050061), and 2-Mercaptoethanol (Gibco #31350010) with Y-27632 dihydrochloride (Tocris Bioscience #1254). After 24 hours, organoids were moved to Ultra-low attachment 6-well plates (Corning #3471) in EM media complemented with Noggin (R&D Systems #6057-NG) and SB 431542 (R&D Systems #1614). Media was changed on day 3 and 4. On day 5, organoids were moved to GFR Matrigel (Corning #354230)-coated 6-well plates in Neural Progenitor Media containing Neurobasal Plus medium (Gibco #A3582901), Glutamax (Gibco #35050061), NEAA (Gibco #11140050), Pen/Strep (Gibco #15140122), N2 supplement (Gibco #17502048), and bFGF (R&D Systems #233-FB). Media was changed on day 6, 8, and 10. On day 12, rosettes were enzymatically lifted with STEMdiff Neural Rosette Selection Reagent (STEMCELL Technologies #05832) and plated NSCs on PLO (Sigma-Aldrich #P4957) /Laminin (Sigma-Aldrich #L2020)-coated plates at a density of 75,000 cells per cm^2^. NSCs were cultured in feeder-free condition with NSC media containing Neurobasal Plus medium (Gibco #A3582901), Glutamax (Gibco #35050061), NEAA (Gibco #11140050), Pen/Strep (Gibco #15140122), N2 supplement (Gibco #17502048), bFGF (R&D Systems #233-FB), B27 supplement (Gibco #17504044), EGF (R&D Systems #236-EG), Insulin (Sigma-Aldrich #I9278), and D-Glucose (Gibco #A2494001).

### NSC differentiation to CO

NSCs were cultured in feeder-free conditions and differentiated into COs through a 3-phase process. In phase 1, on day 0, NSCs were plated in AggreWell 800 plate (STEMCELL Technologies #34811) at density of 3×10^6^ cells per well on an anti-adherence rinsing solution (STEMCELL Technologies #07010)-treated 24-well plate in EB Formation media (STEMCELL Technologies #05893) with Y-27632 dihydrochloride (Tocris Bioscience #1254). On day 1, organoids were moved to Ultra-low attachment 6-well plates (Corning #3471) in NSC media containing Neurobasal Plus medium (Gibco #A3582901), Glutamax (Gibco #35050061), NEAA (Gibco #11140050), Pen/Strep (Gibco #15140122), N2 supplement (Gibco #17502048), bFGF (R&D Systems #233-FB), B27 supplement (Gibco #17504044), EGF (R&D Systems #236-EG), Insulin (Sigma-Aldrich #I9278), and D-Glucose (Gibco #A2494001). Media was changed every other day until day 17. In phase 2, starting day 18, organoids were cultured in Low-cAMP (LC) media containing Neurobasal Plus medium (Gibco #A3582901), cAMP (Sigma-Aldrich #D0627), Ascorbic acid (Tocris Bioscience #4055), BDNF (PeproTech #450-02), NT3 (PeproTech #450-03), GDNF (PeproTech #450-10), Glutamax (Gibco #35050061), NEAA (Gibco #11140050), Pen/Strep (Gibco #15140122), B27 (Gibco #17504001), and N2 supplement (Gibco #17502048). Media was changed every other day until day 45. In phase 3, starting day 46, organoids were cultured in Brainphys+ (BF+) media containing Brainphys basal medium (STEMCELL Technologies #05790), cAMP (Sigma-Aldrich #D0627), ascorbic acid (Tocris Bioscience #4055), BDNF (PeproTech #450-02), GDNF (PeproTech #450-10), Glutamax (Gibco #35050061), NEAA (Gibco #11140050), Pen/Strep (Gibco #15140122), N21-MAX (R&D Systems #AR008), Chemically Defined Lipid (Gibco #11905031), Mouse laminin (Gibco #23017015), and N2 -Plus (R&D Systems #AR003). Media was changed every other day until day 89. Starting day 90, media was changed 2 times per week.

### qPCR

Cells and organoids were cultured in feeder-free conditions. Removed media from culture dish. Rinsed cells and organoids with phosphate-buffered saline (PBS) one time. Lysed cells and organoids with TRIzol Reagent (Invitrogen #15596026) at 4°C for 15 minutes. Extracted and purified RNA from cell lysates using Direct-zol RNA Miniprep kit (Zymo Research #R2052). Generated cDNA from RNA using RevertAid First Strand cDNA Synthesis Kit (Thermo Scientific #K1622). Used 10 ng cDNA per reaction, 5 μ M of primers and PowerUp SYBR Green Master Mix (Applied Biosystems #A46112). Each sample ran three reactions for technical replicates. Reactions were conducted by StepOne Plus Real Time PCR system (Thermo Fisher Scientific). Delta CT was calculated using Human actin as the housekeeping..

### Immunocytochemistry for cryosections

Moved organoids to Ultra-low attachment plates (Corning #3471). Rinsed organoids with PBS one time and removed all liquid. Fixed organoids with 4% PFA (BioWorld #30450002) at room temperature for 20 minutes. Removed all liquid. Incubated organoids with 30% sucrose at 4°C for 48 hours. Moved organoids to cryo-molds (Epredia #58950). Removed all liquid. Embedded organoids with embedding media containing OCT (Sakura Finetek #4583) and 30% sucrose. Froze organoids in -80°C for 24 hours. Cryosectioned organoids in 20-micron thickness and mounted sections on Superfrost Plus Microscope Slides (Fisherbrand #22037246). Washed slides with PBS one time and let dry. Used PAP pen (Vector Laboratories #H-4000) to draw barriers surrounding the sections and let dry. Blocked sections with 10% Normal Goat Serum (Abcam #Ab7481) in 0.1% Triton X-100 at room temperature for 1 hour. Removed all liquid. Incubated sections with primary antibody at 4°C overnight. Removed primary antibody. Washed with PBS three times for 15 minutes each and let dry. Incubated with secondary antibody at room temperature for 1 hour. Removed secondary antibody. Washed with PBS three times for 15 minutes each and let dry. Incubated with Hoechst at room temperature for 10 minutes. Removed Hoechst. Washed with PBS one time for 15 minutes. Used appropriate amount of Fluoromount Aqueous Mounting solution (Sigma-Aldrich #F4680) and covered slides with cover slip (Thomas Scientific #1202F66). Air dried the slides for 30 minutes and sealed the edges with clear nail polish. Let dry for 10 minutes prior to imaging.

### Immunocytochemistry for 2D culture

Cells were cultured in feeder-free conditions. Removed media from culture dish. Rinsed cells with PBS one time. Fixed cells with 4% PFA (BioWorld #30450002) at room temperature for 20 minutes. Removed all liquid. Rinsed cells with TBS Wash Buffer for IHC (Invitrogen #00495456) two times for 2 minutes each. Permeabilized cells with 0.1% Triton at room temperature for 15 minutes. Removed all liquid. Blocked cells with 3% BSA (BioWorld #22070004) in PBS at room temperature for 1 hour. Removed all liquid. Incubated cells with primary antibodies at 4°C overnight. Removed primary antibodies and rinsed cells with wash buffer three times for 5 minutes each. Incubated cells with secondary antibodies at room temperature for 1 hour. Removed all liquid. Rinsed cells with wash buffer three times for 5 minutes each. Incubated cells with Hoechst solution at room temperature for 5 minutes. Rinsed cells with wash buffer for 5 minutes. Added TBS wash buffer to preserve cells.

### RNA sequencing

Cells and organoids were cultured in feeder-free conditions. Removed media from culture dish. Rinsed cells and organoids with phosphate-buffered saline (PBS) one time. Lysed cells and organoids with TRIzol Reagent (Invitrogen #15596026) at 4°C for 15 minutes. Extracted and purified RNA from cell lysates using Direct-zol RNA Miniprep kit (Zymo Research #R2052). Samples were prepared according to Novogene sample submission guidelines and subsequently used to create strand-specific RNA libraries and sequenced on Illumina NovaSeq 6000 (PE150). Sequencing reads were de-multiplexed and aligned to the Hg38 reference genome with STAR Universal Aligner version 2.5.3a using the following settings: Indexed Reference Genome: Ensembl reference genome and annotation files for Hg38 release 77 were downloaded and compiled into a single file, Genome was indexed using the following arguments “STAR -runMode genomeGenerate -runThreadN 12 – genomeDir/STAR INDEX HG38 -genomeFastaFiles GRCh38 r77.all.fa – sjdbGTFfile Homo sapiens.GRCh38.77.gtf – sjdbOverhang 149”; Sample Read Alignment: alignment of each sample’s reads was performed with the following arguments: “STAR - runThreadN 24 -genomeDir/STAR INDEX HG38 -outFileNamePrefix/STAR/SampleName - outSAMtype BAM SortedByCoordinate -outWigType bedGraph -quantMode TranscriptomeSAM GeneCounts -readFilesCommand zcat -readFilesln Sample-R1.fastq.gz Sample-R2.fastq.gz”. Differential Expression (DE) analysis was performed with edgeR software in R Studio. Gene count files were combined into a single file and compared to the respective isogenic cell state. DE genelists from pairwise comparisons were exported into .csv files and utilized for GO term analysis using DAVID. Volcano plots were generated using ggplot2 software in R Studio. Heatmaps were generated using pheatmap software in R Studio. DEG interaction STRING maps were generated using the STRING-database (STRING-db).

### Western blot

Organoids were cultured in feeder-free conditions. Protein samples were collected from organoids after day 70 with at least 3 organoids per sample. Removed media from culture dish. Rinsed organoids with PBS one time. Added 1X HALT (Thermo Scientific #78440) in RIPA (Thermo Scientific #89901) to organoids. Sonicated organoids and centrifuged at 15,000 RCF at 4°C for 15 minutes. Collected supernatant in new Eppendorf tubes. Determined protein concentration using BCA kit (Thermo Scientific #23225). Prepared protein samples with Protein Loading Buffer (National Diagnostics #EC-887) and RIPA (Thermo Scientific #89901). Incubated protein sample mix at 70°C for 10 minutes. Loaded samples into 4-20% Mini-PROTEAN TGX Precast Protein Gels (BioRad #4561094) in running buffer containing 10% SDS (Invitrogen #AM9822) in 1X Tris-Glycine solution (BioRad #1706435). Ran gel at 60V for 20 minutes followed by 120V for 1 to 2 hours. Activated PVDF membrane (BioRad #1620260) using methanol. Transferred gel onto PVDF membrane (BioRad #1620260) in transfer buffer containing 1X Tris-Glycine solution (BioRad #1706435) and 20% Methanol at 90V for 1.5 hours or at 30V for 16 hours. Blocked membrane with 10% Fish Serum Blocking Buffer (Thermo Scientific #37527) at room temperature for 1 hour. Incubated membrane with primary antibodies at 4°C overnight. Removed primary antibodies. Washed membrane with 0.1% TBS (Fisher Bioreagents #BP2471)-Tween (Thermo Scientific #J20605-AP) three times for 5 minutes each. Incubated membrane with secondary antibodies at room temperature for 1 hour. Removed secondary antibodies. Washed membrane with 0.1% TBS (Fisher Bioreagents #BP2471)-Tween (Thermo Scientific #J20605-AP) three times for 5 minutes each. After imaging, stripped membrane with Restore PLUS Western Blot Stripping Buffer (Thermo Scientific #46430) at room temperature for appropriate amount of time until no bands were shown. Blocked membrane in 10% Fish Serum Blocking Buffer (Thermo Scientific #37527) at room temperature for 15 minutes. Incubated membrane with primary antibodies at 4°C overnight. Removed primary antibody. Washed membrane with 0.1% TBS (Fisher Bioreagents #BP2471)-Tween (Thermo Scientific #J20605-AP) three times for 5 minutes each. Incubated membrane with secondary antibodies at room temperature for 1 hour. Removed secondary antibodies. Washed membrane with 0.1% TBS (Fisher Bioreagents #BP2471)-Tween (Thermo Scientific #J20605-AP) three times for 5 minutes each. Imaged with Odyssey CLx (LI-COR) and results were quantified with Empiria Studio.

### Phospho-kinase array

Used Proteome Profiler Human Phospho-Kinase Array Kit (R&D Systems #ARY003C) and followed manufactured guidelines. Briefly, protein from day 123 organoids were loaded at total 200 μ g per sample. Imaged using BioRad ChemiDox XRS+ (BioRad) and results were quantified with ImageJ.BioRad) and results were quantified with ImageJ.

### Plasmid assembly

“PPP2R5D-coOPT” plasmid was cloned using Gibson assembly, inserting codon-optimized PPP2R5D cDNA into the pAAV.CMV.Luc.IRES.EGFP.SV40 (#105533) backbone from Addgene. Plasmids used for mammalian experiments were purified with QIAprep Spin Miniprep Kit (Qiagen, Germantown, MD, USA,#27106) and Plasmid Plus Midi Kit (Qiagen, Germantown, MD, USA,#12945).

### AAV production

HEK293T cells were triple transfected with the transfer plasmid and Helper and Rep/Cap (AAV serotype 2/9) plasmids at approximately 70% confluency. Three days post-transfection, cells were harvested and lysed with lysis buffer. Cell pellets and supernatant were processed with PEG8000 (Sigma) for virus precipitation. For purification of AAVs, a 60% iodixanol gradient was used for ultracentrifugation. Fractions of AAV were removed and resuspended in Lactated Ringer’s Solution, then concentrated by centrifugation with 50 kDa centrifugal concentrator columns (Sartorius, Fremont, CA, #VS2032). Concentrated AAV was then viral genomes (vg) quantified by qPCR using AAVpro Titration Kit (Takara Bio USA, San Jose, CA, #6233) and stored at 4 °C until use.

### AAV transduction in organoids

Organoids were cultured in feeder-free conditions. Moved organoids to a 24-well Ultra-low attachment plate (Corning #3473) with 5 organoids/well. Added 10^11^ vg total AAV to 1 mL of culture media/well. Incubated organoids in AAV media for at least 24 hours. Added 1mL culture media/well and incubated for 24 hours. Fully changed media (48 hours post AAV media) and resumed regular media-change schedule for at least 14 days prior to sample collection.

### High-density MicroElectrode Array

Maxwell MaxTwo system (MaxWell BioSystems) was used. Incubated wells of MaxTwo 6-well plate (MaxWell Systems) in 1% Terg-a-zyme at room temperature for 2 hours. Removed all liquid and wash wells with MilliQ water three times. Autoclaved plate lids prior to proceeding in tissue culture hoods. In tissue culture hoods, incubated plate with 70% ethanol at room temperature for 30 minutes. Removed all liquid and dried plate with vacuum or air dry. Added culture media to wells and incubated plate in incubator overnight. Removed media from wells and wash wells with sterile water three times. Removed all liquid and dry plate with vacuum or air dry. Added 0.07% PEI (Sigma-Aldrich #P3143) to wells and incubated plate in incubator for 1 hour. Removed all liquid and washed wells with sterile water three times. Removed all liquid and dry plate with vacuum or air dry. Added 0.04 mg/mL Laminin (Sigma-Aldrich #L2020) in culture media to wells and incubated plate in incubator overnight. Removed all liquid and dried plate with vacuum or air dry. Moved organoids to wells with one organoid/well. Adjusted position of organoid and allowed maximum surface area of contact with chip. Used Absorption Spears (Fine Science Tools #1810501) to dry the space in between organoid and chip and let organoid settle on chip at room temperature for 5 minutes. Added 50 uL culture media/well and incubated plate in incubator for 2 hours. Added 2 mL culture media/well and incubated plate in incubator for at least overnight. Resumed regular media-change schedule 48 hours post attachment.

Recording sessions were conducted using MaxLab Live software two times per week from day 7 to 28 post attachment with recording parameters, 512x Gain, 1Hz HPF and 5.50 spike detect threshold and comprised three sequential assays. First, an Activity Scan Assay was performed to survey neuronal activity across all 26,400 electrodes (7 configurations, 30 seconds each), providing a comprehensive spatial overview of network activity. Subsequently, a 5-minute Network Scan Assay was performed using 1,020 electrodes selected based on spike amplitude and spatial distribution, enabling targeted recording from regions of high neuronal activity. Finally, a Neuronal Units 9 scan within the Network Assay module was applied, redistributing electrodes in a 3×3 grid configuration to maximize single-unit recording coverage, facilitating post-hoc spike sorting analysis.

HD-MEA data processing was performed using a custom Python pipeline built on SpikeInterface^37^ and Kilosort4^38^. Raw recordings were preprocessed by bandpass filtering between 300 Hz and 3,000 Hz to isolate action potential signals. Spike sorting was subsequently performed using Kilosort4 to identify putative single units on the chip. To ensure data quality, an automated curation step was applied to remove contaminated or misclassified units based on the following criteria: firing rate < 0.01 Hz, median amplitude > −20 µV, refractory period contamination > 0.15, and presence ratio < 0.75 using the quality metrics module of SpikeInterface^37^. Units failing any of these thresholds were excluded from further analysis, ensuring that only well-isolated, physiologically valid neurons were retained for downstream electrophysiological characterization. Spike times from curated units were used to perform network burst analysis using a custom Python algorithm. Network synchrony was computed by tracking co-active units across adaptive time bins, and bursts were detected based on population synchrony peaks exceeding a data-driven threshold, with candidates filtered by neuronal participation, burst density, and firing rate criteria. Detected bursts were characterized by rate, duration, and spikes per burst for cross-condition comparison. Unit-level metrics, including mean firing rate and spike amplitude, as well as waveform metrics such as peak-to-valley time and repolarization slope, were extracted using the quality metrics and template metrics modules within SpikeInterface^37^. All statistical comparisons between conditions were performed using Welch’s independent t-test, with a significance threshold set at p < 0.05.

### Rapamycin treatment for Western b lot

Organoids were cultured in feeder-free conditions. Moved organoids to 24-well Ultra-low attachment plate (Corning #3473) with 5 organoids/well. Added Rapamycin (MedChemExpress #HY-10219) to culture media to a final concentration of 100 nM/well. Added Rapamycin media to organoids. Added 100% DMSO (Sigma-Aldrich #D2650) to culture media to a final concentration of 2%/well. Added DMSO media to organoids as vehicle control. Incubated organoids in incubator for 45 minutes to 1 hour prior to protein collection.

### Rapamycin treatment for MEA

Organoids were attached to MaxTwo plates (MaxWell Systems). For each recording session, recorded b aseline recordings and added Rapamycin (MedChemExpress #HY-10219) to media to a final concentration of 20 nM/well. Incubated plate in incubator for 1 hour and recorded Drug recordings. Recorded using Maxwell MaxTwo system (MaxWell Systems) two times per week from day 7 to 28 post attachment with recording thresholds at 512x Gain, 1 Hz HPF, and 5.50 spike threshold.

---

**Supplement Figure 1. Organoids display cerebral cortical-like nature throughout development.**

(A) Representative 20X images of Healthy, E198K, and E200K COs on day 18, 46, and 90 positive for SATB2, TBR2, CTIP2, TUJ1, MAP2, NeuN, PAX6, vGLUT1, TH, S100B, Nestin, and SOX2. Scale bar represents 50um (20X).

(B) Delta CT of expressions of PAX6, FOXG1, NeuroD1, BDNF, S100B, MAP2, NeuN, vGLUT2, GAD1, TH, and SYN1 in COs on day 18, 46, and 90. One-way ANOVA. * p ≤ 0.05; ** p ≤ 0.01; *** p ≤ 0.001; **** p ≤ 0.0001.

**Supplement Figure 2: Bulk RNA sequencing data reveals significant transcriptomic changes across different stages of mutant organoids.**

(A) Stemness markers enrichment heatmap (left) and astrocyte markers enrichment heatmap (right) of Healthy, E198K, and E200K iPSCs, NSCs, early, mid, and late stages of organoids.

(B) Venn diagrams of overlapping upregulated (top) and downregulated (bottom) DEGs between E198K and E200K organoids in early (left), mid (middle), and late (right) stages.

(C) GO term analyses of upregulated (top) and downregulated (bottom) DEGs of Healthy against mutant organoids in early (left), mid (middle), and late (right) stages.

(D) KEGG pathway analyses of upregulated (top) and downregulated (bottom) DEGs of Healthy against mutant organoids in early (left), mid (middle), and late (right) stages.

**Supplement Figure 3. GABA B R2 is a potential substrate of PP2A-B56.**

(A) Western Blot of Phosph-GABA B R2 S783 and Pan GABA B R2 in Healthy, E198K, and E200K COs with GAPDH housekeeping. Quantification of WB bands by Empiria Studios. One-way ANOVA. * p ≤ 0.05; ** p ≤ 0.01; *** p ≤ 0.001; **** p ≤ 0.0001.

(B) Delta CT of expressions of GABA B R2 in COs on day 86 (no treatment) /17 days post transduction (AAV treated) by qPCR. Two-way ANOVA. * p ≤ 0.05; ** p ≤ 0.01; *** p ≤ 0.001; **** p ≤ 0.0001.

**Supplement Figure 4. Longitudinal network activities in COs are stable across the recording period.**

Line-graphs of quantifications of active area, firing rate, spike amplitude, burst frequency, burst duration, burst peak firing rate, spikes within bursts, number of spikes per burst, and interburst intervals in COs from DOC7 to 28.

**Supplement Figure 5. Overexpression of PPP2R5D in COs through AAV transduction.**

(A) Graphic of structure of AAV with codon-optimized PPP2R5D and eGFP.

(B) Quantification of %GFP by ImageJ. NT %GFP expression 0%, 0.75ug %GFP expression 44%, and 1.5ug %GFP expression 66%.

(C) Western Blot of PPP2R5D in HEK293 cells 72 hours post-transfection with β-tubulin housekeeping. Quantification of WB bands by Empiria Studios.

(D) Representative 10X images of Healthy, E198K, and E200K COs positive for GFP 25 days post transduction. Scale bar represents 300um.

(E) Delta CT of expressions of wildtype *PPP2R5D* and codon-optimized *PPP2R5D* in COs 17 days post transduction by qPCR. Unpaired t test. * p ≤ 0.05; ** p ≤ 0.01; *** p ≤ 0.001; **** p ≤ 0.0001.

(F) Western Blot of PPP2R5D in Healthy, E198K, and E200K Cos 17 days post transduction with GAPDH housekeeping. Unpaired t test. * p ≤ 0.05; ** p ≤ 0.01; *** p ≤ 0.001; **** p ≤ 0.0001.

(G) Representative images of Network Activity plot of Healthy, E198K, and E200K COs 37 days post transduction on Maxwell MaxTwo MEA system.

(H) Quantification of MEA recordings by Maxwell MaxTwo system. Two-way ANOVA. * p ≤ 0.05; ** p ≤ 0.01; *** p ≤ 0.001; **** p ≤ 0.0001.

**Supplement Figure 6. Overexpression of *PPP2R5D* in COs results in downstream transcriptomic, protein functional, and electrophysiological changes.**

(A) Delta CT of expressions of WDFY1 and XBP1 in COs 17 days post transduction by qPCR. Two-way ANOVA. * p ≤ 0.05; ** p ≤ 0.01; *** p ≤ 0.001; **** p ≤ 0.0001.

(B) Phospho-kinase array images of Healthy, E198K, and E200K no treatment and AAV-treated COs.

(C) Quantifications of phospho-kinase arrays by ImageJ.

## Notes

### Competing Interest Statement

The authors have declared no competing interest.

